# Predicting COVID-19 Severity with a Specific Nucleocapsid Antibody plus Disease Risk Factor Score

**DOI:** 10.1101/2020.10.15.341743

**Authors:** S. Sen, E.C. Sanders, K.N. Gabriel, B.M. Miller, H.M. Isoda, G.S. Salcedo, J.E. Garrido, R.P. Dyer, R. Nakajima, A. Jain, A.-M. Caldaruse, A.M. Santos, K. Bhuvan, D.F. Tifrea, J.L. Ricks-Oddie, P.L. Felgner, R.A. Edwards, S. Majumdar, G.A. Weiss

## Abstract

Effective methods for predicting COVID-19 disease trajectories are urgently needed. Here, ELISA and coronavirus antigen microarray (COVAM) analysis mapped antibody epitopes in the plasma of COVID-19 patients (n = 86) experiencing a wide-range of disease states. The experiments identified antibodies to a 21-residue epitope from nucleocapsid (termed Ep9) associated with severe disease, including admission to the ICU, requirement for ventilators, or death. Importantly, anti-Ep9 antibodies can be detected within six days post-symptom onset and sometimes within one day. Furthermore, anti-Ep9 antibodies correlate with various comorbidities and hallmarks of immune hyperactivity. We introduce a simple-to-calculate, disease risk factor score to quantitate each patient’s comorbidities and age. For patients with anti-Ep9 antibodies, scores above 3.0 predict more severe disease outcomes with a 13.42 Likelihood Ratio (96.7% specificity). The results lay the groundwork for a new type of COVID-19 prognostic to allow early identification and triage of high-risk patients. Such information could guide more effective therapeutic intervention.

**Significance statement:** The COVID-19 pandemic has resulted in over two million deaths worldwide. Despite efforts to fight the virus, the disease continues to overwhelm hospitals with severely ill patients. Diagnosis of COVID-19 is readily accomplished through a multitude of reliable testing platforms; however, prognostic prediction remains elusive. To this end, we identified a short epitope from the SARS-CoV-2 nucleocapsid protein and also a disease risk factor score based upon comorbidities and age. The presence of antibodies specifically binding to this epitope plus a score cutoff can predict severe COVID-19 outcomes with 96.7% specificity.

## Introduction

The COVID-19 pandemic has triggered an ongoing global health crisis. More than 108.2 million confirmed cases and 2.3 million deaths have been reported worldwide as of February 16, 2021 (1). The virus that causes COVID-19, severe acute respiratory syndrome coronavirus (SARS-CoV-2), belongs to the same family of viruses responsible for respiratory illness linked to recent epidemics – severe acute respiratory syndrome (SARS-CoV-1 termed SARS here) in 2002-2003 and Middle East respiratory syndrome (MERS) in 2012 (2). The current and previous outbreaks suggest coronaviruses will remain viruses of concern for global health.

Many risk factors and comorbidities, including age, sex, hypertension, diabetes, and obesity, can influence COVID-19 patient outcomes (3). Analysis of patient immune parameters has linked disease severity to elevated levels of biomarkers for inflammation (c-reactive protein and cardiac troponin I), organ damage (aspartate aminotransferase, abbreviated AST, and hypoalbuminemia), immune hyperactivity (IL-6 and IL-10), and clotting (D-dimer) (4). Mortality in COVID-19 is often caused by multi-organ injury and severe pneumonia attributed to an excessive immune response, termed a cytokine storm (5). Given the rapid and wide spectrum of COVID-19 disease progression, a more precise prognostic linking disease risk factors and specific immune responses can potentially predict disease trajectories and guide interventions.

One hypothesis to explain differences in severity of COVID-19 implicates weakly binding, non-neutralizing antibodies (Abs) to SARS-CoV-2 proteins (6). However, the potential harm of these suboptimal Abs in COVID-19 patient outcomes remains ill-defined. Furthermore, a recent review on antibody-dependent enhancement of SARS-CoV-2 stated, “At present, there are no known clinical findings, immunological assays or biomarkers that can differentiate any severe infection from immune-enhanced disease, whether by measuring antibodies, T cells or intrinsic host responses (7).” This conclusion inspired our study.

SARS-CoV-2 encodes four major structural proteins – spike (S), nucleocapsid (N), membrane (M), and envelope (E). The S, N, and M proteins from SARS elicit an Ab-based immune response (8, 9). The Ab response and its effects on disease progression in SARS-CoV-2 remain under investigation (10, 11). Bioinformatics has predicted >55 Ab binding epitope regions from SARS-CoV-2 (12–17). The epitopes for N, M or E proteins are less well-characterized than for S protein. Several studies have reported comprehensive epitope mapping of the antibody response to SARS-CoV-2 (18–21). Here, we sought to characterize epitopes from SARS-CoV-2 and their correlations with disease severity. ELISAs with phage-displayed epitopes (phage ELISAs) and coronavirus antigen microarray (COVAM) analysis (22) examined plasma samples from COVID-19 patients (n = 86). The results demonstrate that Abs to a specific epitope from N protein plus disease risk factors strongly correlate with COVID-19 disease severity.

## Results

### Design and production of candidate epitopes

Twenty-one putative SARS-CoV-2 epitopes were predicted through bioinformatics (12–14) and structure-based analysis. The candidate epitopes span the S, N, M, or E proteins and are on average 34 amino acids in length (**Fig. 1** and **Table S1**). These epitopes were phage-displayed as fragments of the full-length protein and were likely unstructured. Here, epitope refers to the predicted region of the antigenic protein recognized by the antibody’s paratope. The structure of S protein bound to a neutralizing antibody (23, 24) provided the starting point for 12 of these antibody epitopes. Epitopes were designed to potentially isolate even suboptimal Abs binding to small portions of these structural proteins; such suboptimal Abs were hypothesized to provide insight into disease severity. After display of each potential epitope on the surface of phage, the quality of the epitopes was evaluated by PCR, DNA sequencing, and QC ELISA (**Fig. S1**). A total of 18 phage-displayed, putative epitopes passed quality control PCR, and were selected for further study.

**Fig. 1.**
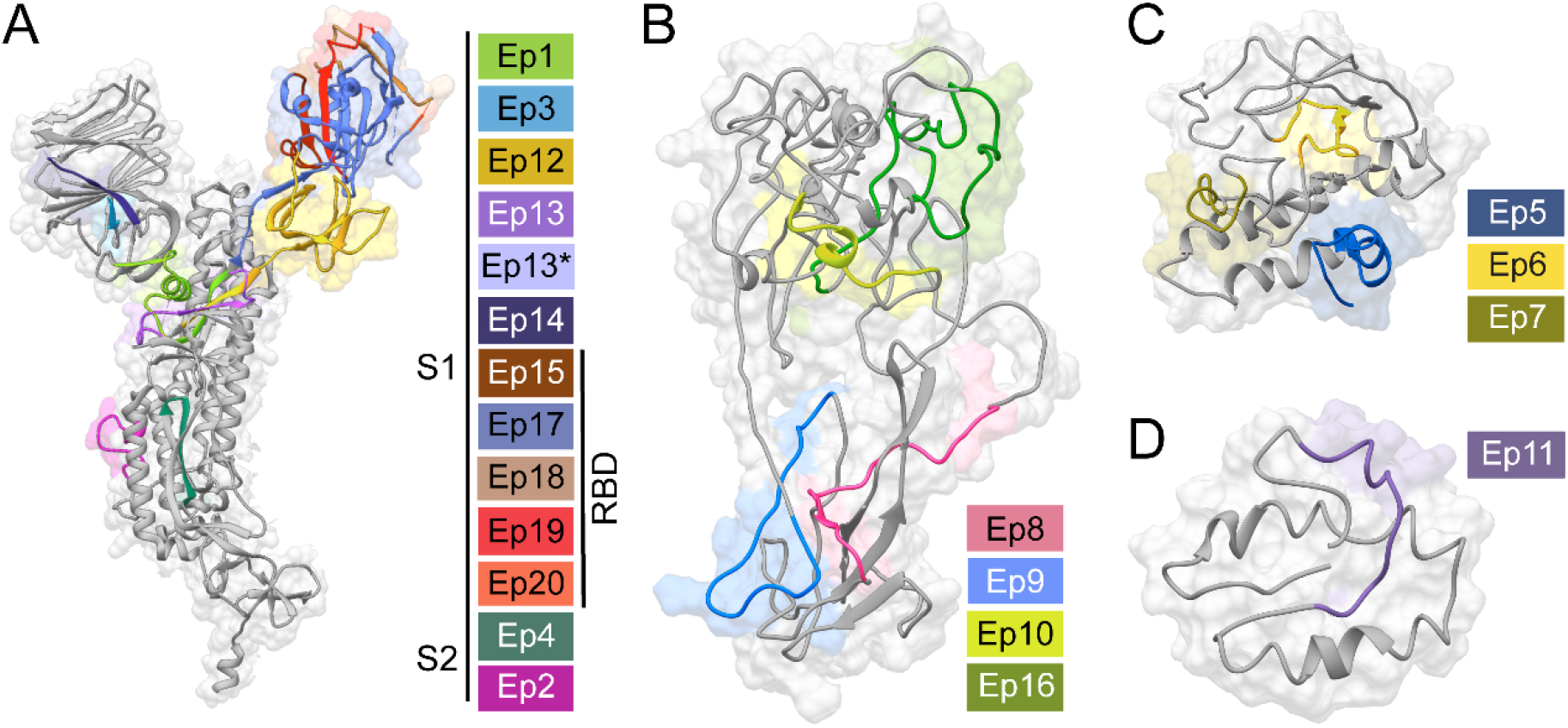
Predicted SARS-CoV-2 epitopes examined by phage ELISA. Structural models (gray) of the SARS-CoV-2 **A**) S, **B**) N, **C**) M, or **D**) E proteins illustrate our epitope design (colored). Sequence Ep13* has the mutation D614G, which increases the fitness of SARS-CoV-2 (25,26). The depicted structural models were derived from an S protein X-ray structure (PDB: 6VXX) (23) or computation modeling of N, M, and E proteins (Protein Gene Bank: QHD43423, QHD43419, and QHD43418, respectively) (27). **Table S1** provides sequences, sources, and rationale for selections.

### Mapping epitope binding to anti-SARS-CoV-2 Abs

Plasma from COVID-19 patients was subjected to ELISAs with the phage-displayed SARS-CoV-2 epitopes **(Fig. 2A)**. Unless otherwise indicated (e.g., healthy controls), plasma refers to samples from PCR-verified, COVID-19 patients. In this initial assay, plasma was pooled, diluted 100-fold, and coated on a microtiter plate (3 pools of n = 5 patients per pool). Nonspecific interactions were blocked (ChonBlock), and phage-displayed epitopes were added for ELISA. The resultant data were normalized by signal from the corresponding negative control (phage without a displayed epitope). Seven candidate epitopes from the pooled patients were further investigated with a larger number of individual patient samples (n = 28) (**Fig. 2B**). The strongest, reproducible binding was observed for three epitopes from M (Ep6), N (Ep9), and S (Ep20) proteins. Additional COVID-19 plasma samples were profiled for binding to these three epitopes (n = 86 total) (**Fig. 2B**).

**Fig. 2.**
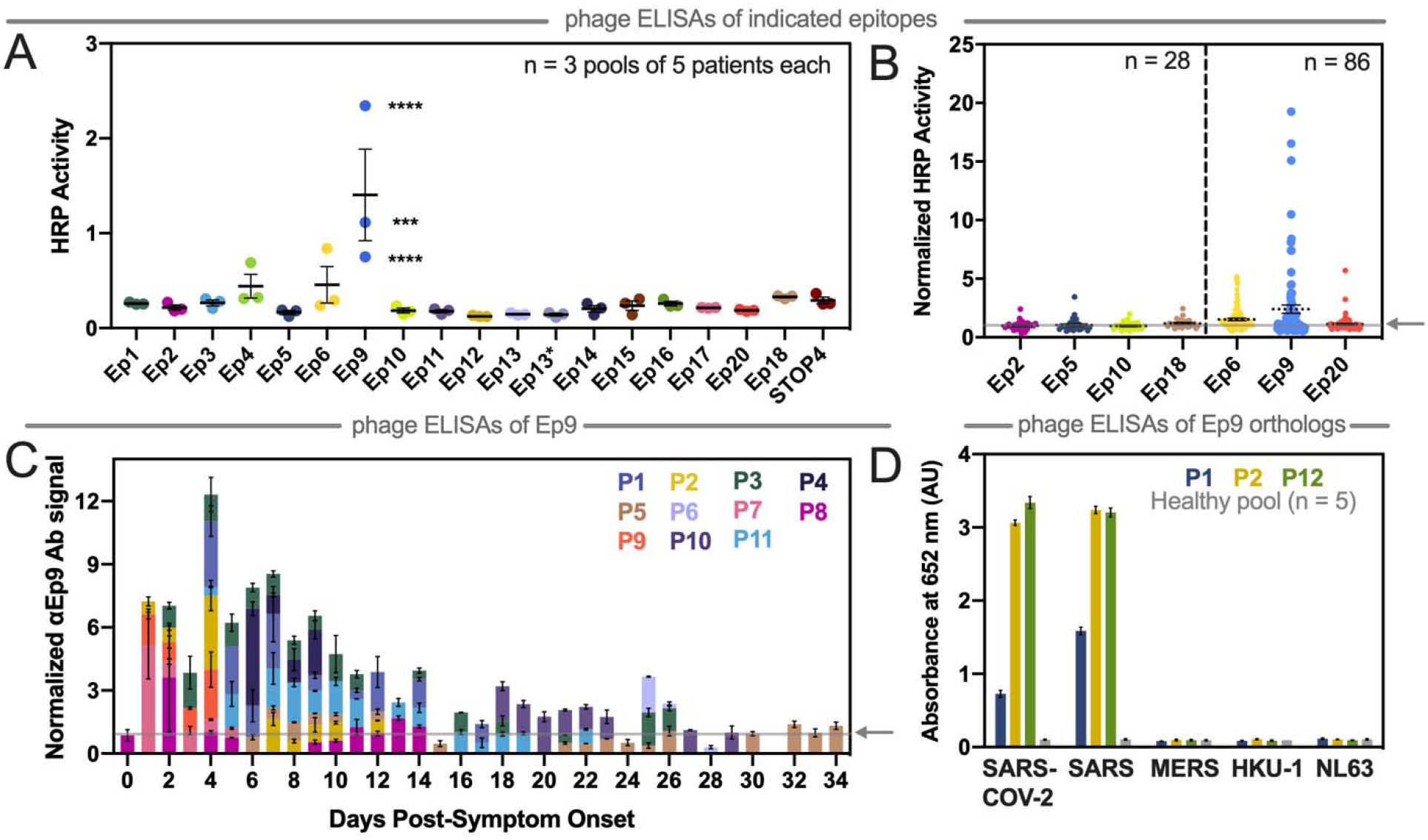
Mapping COVID-19 patient antibody responses with phage-displayed SARS-CoV-2 epitopes. **A)** This phage ELISA with the indicated epitopes (x-axis) examined plasma pooled from patients (n = 3 pools of 5 patients each, 2 technical replicates). STOP4 is the phage negative control. **B)** The epitopes with the highest signals were then further examined by ELISA with plasma from individual patients (n as indicated). **C)** With samples from individual patients (designated as P# and by color) collected at the indicated times, αEp9 Abs were measured. The subset of patients shown here comprise all samples for which longitudinal data was available. **D)** Phage ELISA with samples from patients with strong αEp9 Ab response (two from the longitudinal study and one from the patient population) examines cross-reactive binding to Ep9 or Ep9 orthologs from the indicated coronaviruses (x-axis, 3 technical replicates). The arrow on the y-axis and gray line (panels B and C) represents the negative control used for normalizing the data. Error bars represent SEM (panels A, B, and D) or range of two measurements (panel C).

Only the Ep9 epitope from N protein demonstrated robust, statistically significant antibody binding in 27% of patients (n = 186) (**Figs. 2B** and **S2**). Of these patients, 100 did not have corresponding health information and were not analyzed further in this report. To test non-phage displayed epitopes, dose-dependent binding to Ep9 fused to eGFP (eGFP-Ep9) or to full-length N protein demonstrates that αEp9 IgGs bind its antigen with EC_50_ = 3.22 nM (95% CI = 2.49 to 4.14 nM). This experiment examines plasma samples with the highest IgG response against the N protein in the COVAM assay. Patients without αEp9 Abs have roughly the same level of binding to N protein as observed for αEp9 Abs binding to Ep9. However, such αEp9 Abs appear to add to N protein binding by antibodies; we observe approximately two-fold increase in apparent antibody binding levels for N protein if the patient also has αEp9 Abs (**Fig. S3**). Therefore, the αEp9 response we report cannot solely be due to N protein antigenicity. In patients for whom longitudinal samples were available, the highest levels of αEp9 Abs were observed at days 1 to 14 post-symptom onset (n = 11) and were detectable within 6 days (**Fig. 2C**). In four of these patients, αEp9 Abs persisted after day 14.

### Cross-reactivity of α Ep9 Abs against orthologous epitopes from other coronaviruses

Next, the cross-reactivity of αEp9 Abs was examined with Ep9-orthologs from four phylogenetically related coronaviruses known to infect humans (**Fig. S4A**). Specifically, plasma with αEp9 Abs (n = 3) and pooled plasma from healthy individuals (n = 5) were assayed. The Ep9 epitopes from SARS-CoV-2 and SARS have 90% amino acid sequence homology. Unsurprisingly, this high degree of similarity resulted in a cross-reactive Ep9 epitope, and a strong antibody response was observed to Ep9 epitopes from both viruses (**Fig. 2D**). The coronaviruses, MERS, HKU-1, and NL63 have 52%, 43%, and 8% sequence homology to SARS-CoV-2 Ep9, respectively (**Fig. S4B**). These more distantly related orthologs exhibited no cross-reactivity with the αEp9 Abs. Furthermore, no response was observed to Ep9 in pooled plasma from healthy individuals.

The protein microarray COVAM analysis is a high-throughput serological test for SARS-CoV-2 Ab cross-reactivity with a panel of 61 antigens from 23 strains of 10 respiratory tract infection-causing viruses (22). In this assay, each antigen was printed onto microarrays, probed with human plasma, and analyzed with an ArrayCam imager. COVAM distinguishes between IgG and IgM Abs binding to the full-length N protein (**Fig. S5** and **S6**, respectively). Thus, the COVAM analysis complements the phage ELISA by expanding the scope of antigens surveyed and adding Ab serotype information. The ELISA and COVAM data both demonstrate that αEp9 Abs are highly specific for lineage B betacoronaviruses, and unlikely to be found in patients before their infection with SARS-CoV-2.

### More severe disease and poorer outcomes for α Ep9 patients

Direct comparison of data with full-length N protein from COVAM and Ep9 phage ELISA (n = 40 patients assayed with both techniques) reveals five unique categories of patients (**Fig. 3A**). To enable this comparison, raw data from each assay was normalized as a percentage of the negative control. Category 1 consists of patients without Abs to the N protein. The next categories include patients with IgMs (Category 2) or IgGs (Category 3) binding to N protein, but not Ep9, termed non-Ep9 αN Abs. Category 4 includes patients with αEp9 Abs (both IgMs and IgGs). Category 5 patients have exclusively IgG αEp9 Abs. The αEp9 Abs are only found in patients with IgMs or IgGs against full-length N protein from the COVAM assay; the COVAM analysis thus independently corroborate the phage ELISAs (**Fig. 3A**).

**Fig. 3.**
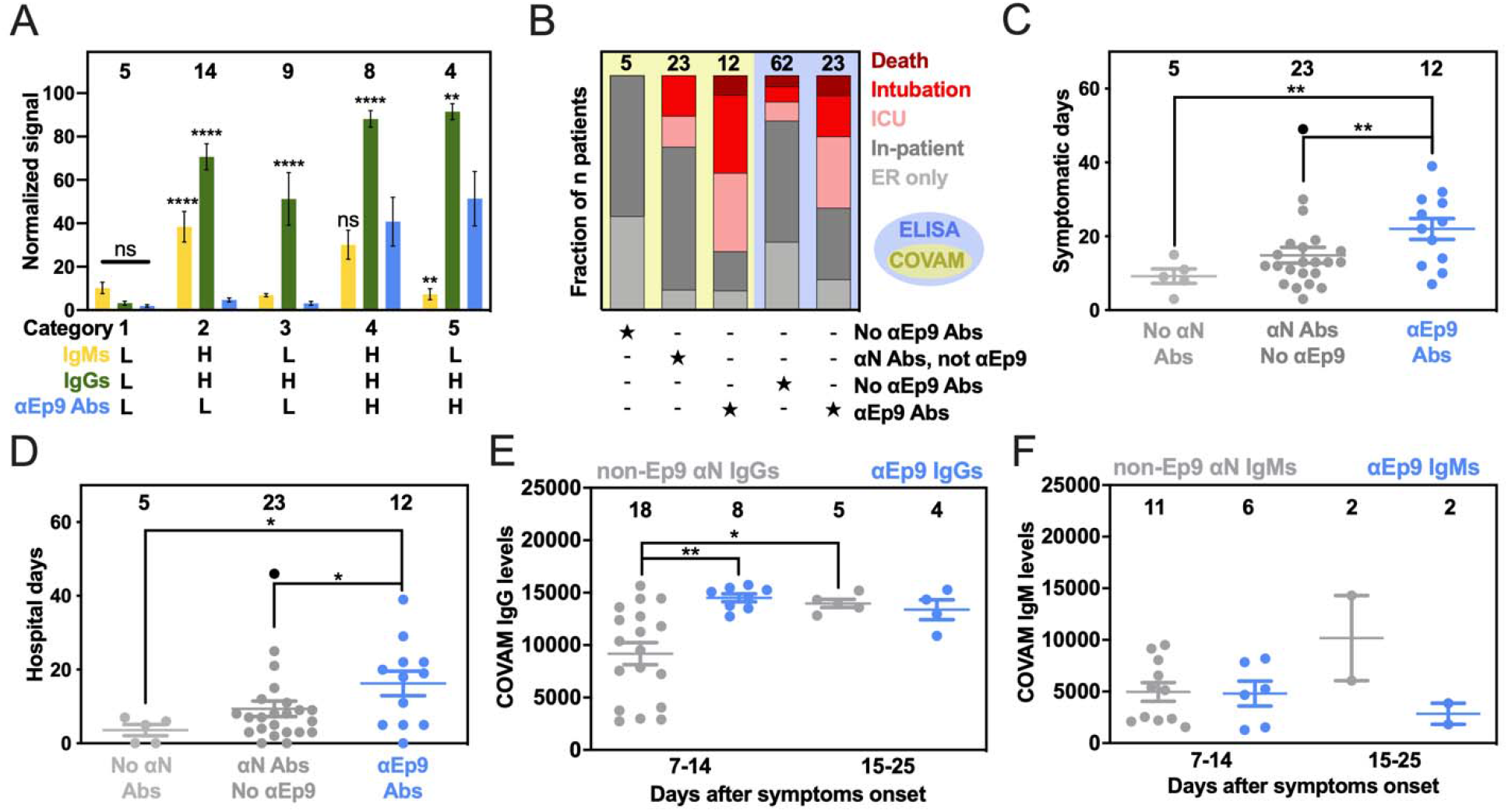
Patients with α Ep9 Abs have more severe disease. **A)** Normalized and categorized data from measurements by COVAM (IgMs in yellow, IgGs in green) and Ep9 phage ELISA (blue). ANOVA comparing COVAM to ELISA with Dunnett’s multiple comparisons yields p-values of **<0.01, ****<0.0001, or ns: not significant. **B)** Disease severity (color) binned by antibody response (COVAM in yellow, or ELISA in blue). Statistical analysis reveals significant differences between distributions of severe and non-severe disease comparing patient categories, p<0.01 (χ^2^) and p<0.001 (Fisher’s exact test) for COVAM and ELISA, respectively. Patients with αEp9 Abs are **C)** symptomatic for longer durations and **D)** spend more days in the hospital than those with other αN Abs or no αN Abs. ANOVA with Tukey’s multiple comparisons yields p-values of *<0.05 and **<0.01. One outlier (black) (ROUT = 0.1%) was omitted from statistical calculations for panels C and D. **E)** The αN IgG appear at high levels early in the course of disease only for αEp9-positive patients, but are lower in non-Ep9, αN-positive patients. After >15 days post symptom onset, αN IgG levels increase for both groups of patients. **F)** However, IgM levels do not change significantly. Error bars depict SEM with the indicated number of patients (n, numbers above columns).

Interestingly, the patients with αEp9 Abs suffer more prolonged illness and worse clinical outcomes compared to patients with non-Ep9 αN Abs or no αN Abs. In this study, severe COVID-19 cases are defined as resulting in death or requiring admission to the ICU or intubation. The fraction of severe COVID-19 cases was 2.5 times higher in αEp9 Abs patients than non-Ep9 αN Abs patients (**Fig. 3B, yellow panel**); the differences in proportions of severe and non-severe αN-positive patients with or without αEp9 Abs are statistically significant (p<0.030, Fisher’s exact test). Patients without αN Abs (Category 1) had less severe symptoms. The αEp9 Abs patients also had longer durations of symptoms and hospital stays relative to non-Ep9 αN Abs and no αN Abs patients (**Figs. 3C** and **D**). A larger data set of patient plasma analyzed by phage ELISA confirmed this conclusion (p<0.0013, Fisher’s exact test) (**Fig. 3B, blue panel**). Our data further demonstrates that asymptomatic COVID-19 patients (n = 3) also tested negative for αEp9 Abs (**Table S2**). The data also reveals early seroconversion of αEp9 IgGs (**Fig. 3E**), but not αEp9 IgMs (**Fig. 3F**).

### Strong correlation between disease severity and comorbidities in patients with α Ep9 Abs

We compared risk factors, clinical parameters, and disease outcomes among patients with αEp9 Abs (n = 23) (**Figs. 4A** and **S7**). A *disease risk factor score* (DRFS) was developed to evaluate the relationship between clinical preconditions and disease severity in patients with αEp9 Abs. The DRFS quantifies a patient’s age, sex, and pre-existing health conditions associated with COVID-19 disease severity and mortality. Risk factors include hypertension, diabetes, obesity, cancer, and chronic conditions of the following: cardiac, cerebrovascular, kidney, and pulmonary (28–31). Using the *age score* from the Charlson Comorbidity Index (32) yields a patient’s DRFS as:

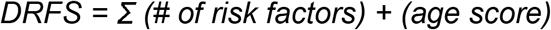

where each risk factor is valued as either 0 or 1 if absent or present, respectively. The DRFS of patients with αEp9 Abs strongly correlates with COVID-19 disease severity (Pearson’s r = 0.72, p-value <0.0001, and R^2^= 0.52) (**Fig. 4A**). The correlation in patients without αEp9 Abs is weak (r = 0.30, p-value = 0.089, R^2^= 0.018) (**Fig. 4A**). Amongst patients with αEp9 Abs (n = 23), a DRFS ≥3 can determine disease severity with 92.3% sensitivity (1/13 false negatives) and 80% specificity (2/10 false positives) (**Fig. 4B**). In the entire study cohort (n = 86), patients with αEp9 Abs and DRFS ≥3 (n = 11) have severe disease with a high degree of specificity (96.7%) and a sensitivity of 44%. Notably, DRFS predicts disease severity only for patients with αEp9 Abs (n = 23), and patients without such Abs (n = 63) had no correlation with disease outcomes.

**Fig. 4.**
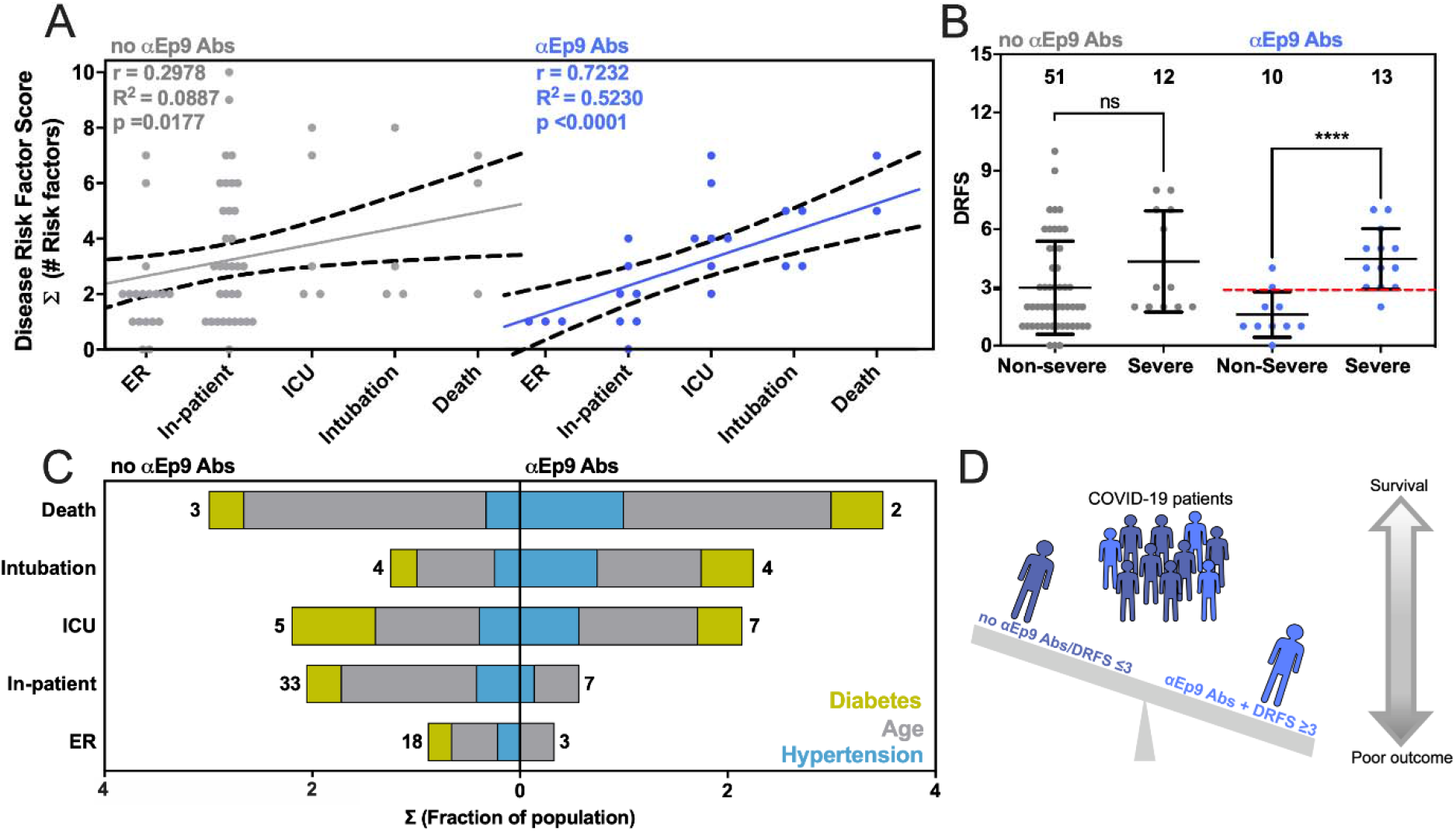
Correlation between disease severity and risk factors in patients with α Ep9 Abs. **A)** The relationship between DRFS and disease severity of COVID-19 patients with αEp9 Abs (blue) or no αEp9 Abs (gray). Each data point represents one patient. The solid lines indicate linear regression fits with 95% confidence intervals (dotted lines), and Pearson’s r-value as noted. **B)** Correlation of disease severity with DRFS in patients with αEp9 Abs. The data depicts a significant correlation between DRFS and disease severity in patients with αEp9 Abs (blue), but not in patients lacking αEp Abs (gray). In αEp9 patients, a DRFS threshold of 3.0 can predict severe disease (red). Two-tailed, parametric t-tests were conducted to compare non-severe and severe disease outcomes of patients with and without αEp9 Abs, where ****p<0.0001. The error bars represent SD with the indicated n. **C)** The color-indicated risk factors (diabetes, hypertension, and age score) are depicted on the x-axis as the fractions of patients in each disease severity category (y-axis). Numbers indicate total patients (n) without αEp9 Abs (left) or with αEp9 Abs (right). The prevalence of risk factors (colors) increases with disease severity in patients with αEp9 Abs, but not in patients without these Abs. **D)** Patients with αEp9 Abs and DRFS ≥3 are predisposed to increased COVID-19 severity and poorer outcomes.

Examining key contributors to high DRFS, the presence of αEp9 Abs correlates with more severe disease in patients who have hypertension, diabetes, or age >50 years. Such correlation is not observed for patients lacking αEp9 Abs (**Fig. 4C**). Such risk factors are prevalent at roughly the same percentages in both populations of patients (**Table S2**). Thus, these risk factors are particularly acute for patients with αEp9 Abs.

### High levels of inflammatory cytokine and tissue damage markers in patients with α Ep9 Abs

COVID-19 patients can have elevated serum concentrations of >20 inflammatory cytokines and chemokines (33). However, information on the cytokine levels and the association with tissue damage and worse COVID-19 outcomes have been inconsistent (33–35). For patients with IL-6 concentrations measured in plasma, patients with (n = 8) or without (n = 11) αEp9 Abs were compared. Interestingly, the comparison uncovered a strong positive sigmoidal association between IL-6 and AST unique to patients with αEp9 Abs (R^2^ = 0.968, Spearman’s r = 1.0, p-value <0.0001, n = 8) (Blue line, **Fig. S8A**); correlation of IL-6 and AST in patients with αEp9 Abs remains strong even after removal of the data point at the highest IL-6 concentration. Conversely, a slight negative trend is observed in patients lacking αEp9 Abs (Spearman’s r = −0.575, p-value= 0.0612, n = 13). Thus, the presence of αEp9 Abs can disambiguate the sometimes contradictory association of IL-6 with disease severity.

## Discussion

This study introduces a two-step test as a prognostic for predicting COVID-19 disease severity and its worst outcomes. Specifically, αEp9 Abs can effectively predict severe disease (specificity 83.6%). However, combining presence of αEp9 Abs with DRFS ≥3 provides much higher specificity (96.7%) for predicting severe disease. Previously, αN IgGs have been recognized as a focal site for an antibody response (18, 19, 21, 36) and associated with disease severity and poor outcomes (11, 36, 37).

The present investigation expands on previous reports that recognize various regions of the RNA binding domain of N protein as focal sites for anti-SARS-CoV-2 antibody response. For example, the phage display-based VirScan identified an epitope region spanning residues 141-196 and microarrays further isolated peptides including residues 134-171, 155-171, 153-190, and 153-171 (18, 19, 21). The above investigations, however, do not find correlations between any these epitopes and disease severity. Our results are confirmed by observations from a patient cohort in Singapore, which identify an epitope (residues 153-170) very similar to Ep9 (residues 152-172) and shows a correlation between antibody response against the epitope and pneumonia and the tissue damage markers (CRP and LDH) (20). In our investigation, we examine in-depth patient clinical histories, test results, disease outcomes ranging from asymptomatic to fatal, and longer longitudinal profiling post-symptom onset, to determine the association of a larger subset of markers and risk factors. Such data allows calculation of the DRFS. Together with the presence of αEp9 Abs, patient DRFS allows early discrimination of severe from non-severe disease outcomes. Additionally, fine epitope mapping demonstrates that αEp9 Abs strongly and uniquely correlate with COVID-19 disease severity relative to other αN Abs.

We hypothesize that the underlying mechanism relating αEp9 Abs to increased disease severity involves an overzealous immune response. Specifically, we observe early seroconversion and strong early upregulation of αEp9 IgGs (**Fig. 3E**). Similar IgG observations have been correlated with poor viral neutralization and clearance, resulting in increased COVID-19 severity (10, 37, 38). Also, high levels of IL-6 are observed for αEp9-positive patients with increased levels of the tissue damage marker AST; this correlation does not exist for patients lacking αEp9 Abs (**Fig. S8A**). The sensitivity to IL-6 concentration before AST-monitored organ damage suggests anti-IL-6 therapeutics could be an effective management in the early and rapidly progressive stages of respiratory distress for αEp9-positive patients (33, 39–43). Since binding to N protein by αEp9 antibodies is unlikely to enhance uptake of SARS-CoV-2, an antibody-dependent enhancement mechanism could invoke antigen uptake by macrophages. This mechanism could stimulate complement activation and the cytokine storm observed here as elevated IL-6 response. Further investigation is required to determine the basis for increased disease severity in αEp9 patients.

The data demonstrate that αEp9 positive patients with DRFS ≥3 are 13.42 times (Likelihood Ratio) more likely to have severe COVID-19 disease symptoms within the study cohort (n = 86). The presence of αEp9 without DRFS is less effective as a prognostic (Likelihood Ratio of 3.17). Despite its high specificity (96.7%), the sensitivity of this two-step test is 44% (n = 86). However, this test could predict a subset of patients with a specific immune response (i.e., early IgG response and IL-6 dependent immune hyperactivity), and could suggest targeted treatment options (e.g., targeting IL-6 and its pathways).

Importantly, αEp9 Abs appear early in the course of disease. Thus, such a prognostic could outperform traditional markers for the cytokine storm such as IL-6, which appears 6-8 days after symptom onset (33, 41); all plasma collected from αEp9 positive patients (n = 7, **Fig. 2C**) between 1 to 6 days post-symptoms onset demonstrate detectable levels of αEp9 IgG (≥ 2 fold over negative control). Early detection of αEp9 Abs in patients could be used to triage and treat COVID-19 prior to the onset of its most severe symptoms; delayed treatments of IL-6 targeting drugs can decrease their efficacy or be counterproductive (33, 39–44) (**Fig. S8B**). The αEp9 Ab biomarker could identify patients most likely to benefit from anti-IL-6 therapeutics and avoid ineffective treatments.

This study demonstrates the usefulness of fine epitope mapping, but the following limitations should be noted. Short linear epitopes, unlike conformational epitopes in larger domains, might not resemble the tertiary structure of an antigen. Post-translational modifications, such as glycosylation were omitted for the phage-displayed S protein epitopes; the COVAM antigens, however, are produced in baculovirus or HEK-293 cells, which could glycosylate the antigens. Our analysis is largely based upon a population of 86 COVID-19 patients and 5 healthy individuals, with the majority of Hispanic descent. The conclusions could be further strengthened with follow-up investigations in a larger population. Additionally, the population examined here only included three asymptomatic individuals, and additional testing is required to verify absence of αEp9 Abs in such patients. The sample size of patients with multiple antibody targets was too limited to allow correlation analysis; future investigations could examine associations between αEp9 and other Abs. Abs recognizing other SARS-CoV-2 structural proteins could also exhibit similar characteristics to αEp9 Abs.

Existing diagnostic platforms could readily be adapted to test for αEp9 Abs, and the DRFS calculation is quite simple to implement (e.g., assay with eGFP-Ep9 fusion demonstrated here). As shown here, αEp9 Abs do not recognize orthologous sequences from closely related coronaviruses, providing good specificity for αEp9 as a prognostic. Previous studies have shown that the high homology of N protein among related coronaviruses can lead to high false positive rates in serodiagnostics with full-length N antigen (45). Thus, the two-step prognostic reported here could mitigate the worst outcomes of COVID-19, particularly for patients at high risk.

## Materials and Methods

Detailed materials and methods for cloning, phage purification, patient sample collection, plasma phage-antibody ELISA, serum COVAM, and statistical analysis are described in the *SI Appendix.*

## Acknowledgements

We thank Dr. Hung Fan and Dr. Donald Forthal for helpful conversations and the patients who donated samples. We gratefully acknowledge the support of the UCI COVID-19 Basic, Translational and Clinical Research Fund (CRAFT), the Allergan Foundation, and UCOP Emergency COVID-19 Research Seed Funding. S.S. was supported by a Public Impact Fellowship from the UCI Graduate Division. K.N.G. was supported by a National Science Foundation Graduate Research Fellowship Program (DGE-1839285). G.S.S and A.M.S thank the Minority Access to Research Careers (MARC) Program, funded by the NIH (GM-69337). J.L.R. was supported by the National Center for Research Resources and the National Center for Advancing Translational Sciences from the NIH (TR001414). D.F.T and R.A.E. were supported by the Experimental Tissue Resource, funded by the Chao Family NCI-Comprehensive Cancer Center Support Grant from the NCI (P30CA062203).

## Abbreviations

COVID-19: Coronavirus disease 2019
ELISA: enzyme-linked immunosorbent assay
COVAM: coronavirus antigen microarray
Ep9: epitope 9
ICU: intensive care unit
SARS-CoV-2: severe acute respiratory syndrome coronavirus
SARS-CoV-1: severe acute respiratory syndrome
MERS: Middle East respiratory syndrome
AST: aspartate aminotransferase
IL-6: interleukin 6
IL-10: interleukin 10
Abs: antibodies
S protein: SARS-CoV-2 spike glycoprotein
N protein: SARS-CoV-2 nucleocapsid protein
M protein: SARS-CoV-2 membrane protein
E protein: SARS-CoV-2 envelope protein
Phage ELISA: ELISA with phage-displayed epitopes
PCR: polymerase chain reaction
QC ELISA: quality control ELISA
PDB: protein data bank
HKU-1: human coronavirus HKU1
NL63: human coronavirus NL63
IgG: immunoglobulin G
IgM: immunoglobulin M
αEp9 Abs: anti-Ep9 antibodies
ANOVA: analysis of variance
DRFS: disease risk factor score

## SI Appendix

General methods and additional experimental data can be found in the *SI Appendix.*

Figs. S1 to S8

Tables S1 to S4

References (46–53)

