## Supplemental Information for "Predicting COVID-19 Severity with a Specific Nucleocapsid Antibody plus Disease Risk Factor Score"

### SI Appendix

#### Materials and Methods

Cloning

For phage display of epitopes, the pm1165a phagemid vector as previously described (*46*) as engineered to encode an N-terminal FLAG-tag and a C-terminal fusion to the P8 coat protein of M13-phage. This template, termed FlagTemplate, was used for subcloning of SARS-CoV-2, SARS, MERS, HKU-1 and NL63 epitopes. A vector map of the FlagTemplate (**Fig. S9)**, cloning procedures, and a list of oligos (**Table S3**) for Q5 site-directed mutagenesis and Gibson Assembly are provided here.

Short (approximately 30 amino acids) putative epitopes for phage display and *E. coli* expression as eGFP fusion peptides in the pet28 vector were cloned via Q5 site-directed mutagenesis according to the manufacturer's instructions. A vector map of Ep9 fused peptide to eGFP, termed eGFP-Ep9, is shown below (**Fig. S10**). For large epitopes (>500 bp), such as Ep17, Gibson Assembly (New England Biolabs) was conducted in two PCR steps with the **FlagTemplate** or pCAGGS containing the SARS-CoV-2 S protein gene (BEI Resources) to generate the vectors and inserts, respectively. The Gibson Assembly (2 µL) or KLD (Kinase, Ligase, DpnI) mix (5 µL) was transformed into Nova Blue *E. coli* competent cells, and transformants were plated on a carbenicillin-supplemented (50 µg/mL) agar plate before incubation at 37 °C overnight. Five single colonies were selected to inoculate 4 mL of SOC media in a 15 mL culture tube supplemented with carbenicillin (50 µg/mL). The seed cultures were incubated at 37 °C with shaking at 225 rpm for 8-12 h. Phagemid DNA was isolated using the QIAprep spin miniprep kit according to the manufacturer's instructions. The successful subcloning of the ORF encoding each epitope was verified via DNA sequencing (Genewiz). The full-length N protein in a pLVX-EF1α-IRES-Puro plasmid was a generous gift from Prof. Rachel Martin (UCI).

Purification and preparation of phage

Phage were propagated and purified using procedures previously described (*47*) with the following changes. A single colony was selected to inoculate 15 mL of 2YT and shaken at 37 °C until the $OD_{600}$ reached 0.6. After incubation at 37 °C for 45 min, 8 mL of the primary culture was used to inoculate 300 mL of 2YT supplemented with carbenicillin (50 µg/mL), kanamycin (20 µg/mL), and isopropyl β-D thiogalactopyranoside (IPTG, 30 µM).

To precipitate the phage, the cultures were centrifuged at 10 krpm (15300 x g) for 10 min at 4 °C. The supernatant was decanted into a centrifuge tube containing 60 mL PEG-8000 (20%, w/v) and NaCl (2.5 M). The tube was inverted 10 times and stored on ice for 30 min followed by an additional centrifugation at 10 krpm (15300 x g) for 20 min at 4 °C. The supernatant was decanted, and tubes were centrifuged for an additional 4 min at 4 krpm (2429 x g) at 4 °C. The pellets were resuspended in PBS (10 mM phosphate, 137 mM NaCl, pH 7.2) with TWEEN 20 (0.05%, v/v) and glycerol (10%, v/v), separated into 1 mL aliquots, flash frozen with liquid nitrogen, and stored at -80 °C. For binding assays via ELISA, the purified phage was thawed on ice, precipitated a second time as before. The quality of each phage preparation was routinely checked by quality control ELISA, termed QC ELISA, to a FLAG peptide fused to the N-terminus of each epitope (**Fig. S9**); additionally, PCR using Oligo69 and Oligo70 followed by DNA sequencing (Genewiz) was performed for every phage preparation. Such quality control allowed for identification of toxic clones; for example, C8, was apparently toxic to *E. coli*, and three protein epitopes failed to express in *E. coli* for unknown reasons. The phage concentration was determined by absorbance at 260 nm using a coefficient of molar absorptivity of 0.003 $nM^{-1}$ $cm^{-1}$ and diluted to 40 nM in PBS.

Expression and Purification of eGFP-Ep9 and N protein

A pET28c plasmid containing Ep9 fused to an N-terminal eGFP (**Fig. S10**) was transformed into BL21 DE3* *E. coli* heat shock, competent cells. A single colony was transferred to LB media (20 mL) supplemented with kanamycin (40 µg/mL) and incubated at 37 °C for 18 h. An aliquot of the starter culture (2.5 mL) was transferred to LB media with 1% glucose (250 mL LB in a 1 L baffled flask). After reaching an $OD_{600}$ between 0.4-0.6, the culture was induced through addition of IPTG (0.5 mM) before incubation at 25 °C for 18 h. The cells were centrifuged (15,300 x g) for 20 min at 4 °C, and the cell pellet was resuspended in lysis buffer (25 mM Tris-HCl and 200 mM NaCl, pH 8.0 and supplemented with protease inhibitor cocktail) followed by sonication. The lysate was subjected to centrifugation (26,892 rcf, 45 min, 4 °C). The supernatant was incubated with charged Ni-IMAC resin overnight on a rotary shaker (150 rpm at 4 °C). The resin was equilibrated in a column, washed with wash buffer (20 mM imidazole in lysis buffer), and the purified protein was eluted using elution buffer (250 mM imidazole in lysis buffer). Elutions containing the purified protein were visualized using 10% or 12% SDS-PAGE (Bio-rad Mini-PROTEAN Tetra electrophoresis system) stained with Coomassie brilliant blue stain (**Fig. S11**). The eluted fractions containing the purified eGFP-Ep9 were pooled and buffer exchanged for 3 column volumes (20 mL) with lysis buffer without imidazole using a 10 kDa cutoff microconcentrator (Vivaspin, Fisher Scientific). The protein concentration was determined by a bicinchoninic acid (BCA) assay or Bradford assay using the estimated MW (http://www.expasy.org). Similar to eGFP-Ep9, the full-length N protein was expressed in 250 ml LB with 1% glucose and induced with 0.25 mM IPTG at $OD_{600}$ = 0.8. Protein overexpression cultures were incubated at 16 °C for 22 h. Lysis and purification were conducted as described above, using N protein lysis buffer (20 mM Tris-HCl, 300 mM NaCl, 5 mM $MgCl_2$, 5 mM BME, 10% glycerol pH 8.0). The purified full-length N protein was analyzed using 10% or 12% SDS-PAGE (**Fig. S11**).

Patient sample collection

The UC Irvine Experimental Tissue Resource operates under a blanket IRB protocol (UCI #2012-8716) that gives ETR personnel 'Honest Broker' status and enables the collection of any fluid or tissue remnant in excess to that needed for clinical diagnosis and distribution to investigators under the conditions of their own IRB approval. Patients undergoing COVID testing in the Emergency Department or on the inpatient service with confirmed COVID (+) pharyngeal swabs, were followed for their blood collections daily. Specimens collected originally for diagnostic purposes were processed and stored by the hospital laboratory in a manner compliant with College of American Pathologists (CAP) standards. EDTA-anticoagulated whole blood was stored for 2 days at 4 °C after clinical diagnosis and released for research purposes. Plasma from heparin-anticoagulated blood was centrifuged immediately after collection and preserved at 4 °C for 3-4 days before being released for research use. All COVID (+) specimens were handled under BSL-2 conditions, aliquoted into screw cap cryovials, and stored at -80 °C long term with constant temperature monitoring. Specimens were coded by the ETR with unique de-identifiers, and accompanying clinical information was stripped of PHI, such that investigators could receive specimens under a Non Human Subjects Determination exemption from the UC Irvine IRB. All samples from SARS-CoV-2 infected patients were inactivated by incubation in a water bath at 56 °C for 30 min (*48*), aliquoted (40 µL each), and stored at -80 °C.

Phage ELISA with plasma

The phage-displayed SARS-CoV-2 epitopes were used in phage ELISAs with patient plasma samples diluted 100-fold in coating buffer (50 mM $Na_2CO_3$, pH 9.6). After incubation in a 96-well Nunc MaxiSorp flat-bottom microtiter plate with shaking at 150 rpm at 4 °C for 12-18 h, plasma was aspirated by a plate washer (BioTek). Next, the plate was treated with 100 µL per well of ChonBlock Blocking/Sample Dilution Buffer (Chondrex, Inc.) for 1 h with shaking at 150 rpm at room temperature and washed three times with wash buffer (0.05% v/v Tween-20 in PBS). The epitope displaying phage and controls were diluted to 1 nM in ChonBlock Blocking/Sample Dilution Buffer and 100 µL were added to each well before incubating for 2 h with shaking (150 rpm) at room temperature. The plate was then washed three times with wash buffer. The primary antibody, anti-M13-HRP (Creative Diagnostics), was diluted 1:5000 in ChonBlock Secondary Antibody Buffer and 100 µL was added per well; the plate was incubated for 1 h at 150 rpm and room temperature. Following three washes with wash buffer, 1-Step Ultra TMB-ELISA Substrate Solution (100 µL per well, ThermoScientific) was added. Absorbance of TMB substrate was measured twice at 652 nm by UV-Vis plate reader (BioTek) after 5 and 15 min of incubation.

##### ELISA of eGFP-Ep9 and full-length N protein with plasma

Varying doses, with a maximum concentration of 1.7 µM, of eGFP-Ep9, eGFP-FLAG or full-length N protein (fl-N) were diluted in PBS pH 8.0, and then immobilized on a 96-well Nunc MaxiSorp flat-bottom microtiter plate before incubatation on a shaker (150 rpm) at 4 °C for 12 to 18 h. After incubation, unattached proteins were removed through aspiratation using a plate washer (BioTek) and wells were blocked with 100 µL ChonBlock Blocking/Sample Dilution Buffer (Chondrex, Inc.) for 30 min with shaking (150 rpm) at room temperature. The plate was then washed three times with wash buffer (0.05% v/v Tween-20 in PBS). Pooled plasma from five patients within each experimental group was diluted 100-fold in ChonBlock Blocking/Sample Dilution Buffer and 100 µL was added to each well before incubating for 1 h with shaking (150 rpm) at room temperature. The plate was then washed three times with wash buffer. The detection antibody, IgG Fc Goat anti-Human, HRP (Invitrogen), was diluted 1:5000 in ChonBlock Secondary Antibody Buffer and 100 µL was added per well; the plate was incubated for 30 min at 150 rpm and room temperature. Following six washes with wash buffer, 1-Step Ultra TMB-ELISA Substrate Solution (100 µL per well, ThermoScientific) was added. Absorbance of TMB substrate was measured twice at 652 nm by UV-Vis plate reader (BioTek) after 5 and 15 min of incubation.

##### Serum coronavirus antigen microarray (COVAM)

COVAM included 61 antigens across respiratory virus subtypes including 11 antigens from SARS-CoV-2 expressed in either baculovirus or HEK-293 cells as previously detailed (**Table S4**) (*22*). These antigens were provided by Sino Biological U.S. Inc. as either catalog products or custom synthesis service products. The antigens were printed onto microarrays, probed with human sera, and analyzed as previously described (*49-51*). Briefly, lyophilized antigens were reconstituted with sterile water to a concentration of 0.1 mg/mL protein in PBS, and printing buffer was added. Antigens were then printed onto ONCYTE AVID nitrocellulose-coated slides (Grace Bio-Labs) using an OmniGrid 100 microarray printer (GeneMachines). The microarray slides were probed with human sera diluted 1:100 in 1X Protein Array Blocking Buffer (GVS Life Sciences, Sanford, ME) overnight at 4°C and washed with TTBS buffer (20 mM Tris-HCl, 150 mM NaCl, 0.05% Tween-20 in ddH2O adjusted to pH 7.5 and filtered) three times for 5 min each. A mixture of human IgG and IgM secondary antibodies conjugated to quantum dot fluorophores Q800 and Q585 respectively was applied to each of the microarray pads and incubated for 2 h at room temperature, and pads were then washed with TTBS three times for 5 min each and dried. The slides were imaged using an ArrayCam imager (Grace Bio-Labs) to measure background-subtracted median spot fluorescence. Non-specific binding of secondary antibodies was subtracted using a saline control. The mean fluorescence of the 4 replicate spots for each antigen was used for analysis.

##### Statistical analysis

The ELISA data were analyzed in GraphPad Prism 8. Since the total antibody content differs from person to person, the raw absorbance values for every patient sample were normalized and represented as the ratio as compared to a negative control. Analysis of variance (ANOVA) with Dunnett's multiple comparisons was performed to determine if values were statistically significant. Correlations between COVAM IgG/IgM and ELISA were determined by plotting normalized values on an XY graph and performing a non-parametric correlation analysis using a Spearman's rank correlation coefficient test.

For data visualization of clinical patient data, trends in data were evaluated using Knime Analytics Platform software. GraphPad Prism was used to calculate column statistics including mean, standard deviation, SEM, p-values, Odds Ratios, and Likelihood Ratios defined as sensitivity / (1 - specificity). ANOVA with Tukey's multiple comparisons test was used to evaluate antibody response and disease severity between patients with αEp9 Abs, non-Ep9, αN Abs, or non αN Abs. Comparisons of patients with αEp9 Abs and non-αEp9 Abs were conducted using unpaired, two-tailed, parametric t-tests. Contingency graphs were statistically evaluated using Fisher's exact test, for groups with binary categorization, and Chi-squared test for groups with multiple categories. Different datasets were fitted with linear or non-linear regression methods, the fit with the higher $R^2$ value was chosen. Correlations between two clinical parameters (e.g., IL-6 and AST) were evaluated using the Pearson coefficient or Spearman coefficients (r) for linear or non-linear regressions, respectively; r-values between 1.0-0.7 were considered strong correlations, r-values between 0.7 and 0.5 were considered a moderate correlation, and values below 0.5 were considered a weak correlation (*52*). The significance of the correlation was evaluated based on p-value <0.05.

#### Supplementary Text

**Table S1. Phage-displayed putative epitopes of SARS-CoV-2 and Ep9 orthologous sequences from SARS, MERS, HKU-1, and NL63.**

| Epitope | Virus | Protein | Residues* | Amino Acid Sequence | Sequence Length | Rationale for selection | Ref. for rationale |
|---|---|---|---|---|---|---|---|
| Ep1 | SARS-CoV-2 | S | 287-317 | DAVDCALDPLSETKCTLKSFTVEKGIYQTSN | 31 | Predicted B-cell epitope in SARS-Cov-2 identified by bioinformatics to have immunodominant region | (*13*) |
| Ep2 | SARS-CoV-2 | S | 802-819 | FSQILPDPSKPSKRSFIE | 18 | Predicted B-cell epitope in SARS-Cov-2 identified by bioinformatics to have immunodominant region | (*13*) |
| Ep3 | SARS-CoV-2 | S | 15-30 | CVNLTTRTQLPPAYTN | 16 | Predicted B-cell epitope in SARS-CoV-2 identified by immuninformatics | (*14*) |
| Ep4 | SARS-CoV-2 | S | 1056-1070 | APHGVVFLHVTYVPA | 15 | Predicted B-cell epitope in SARS-CoV-2 identified by structural biology and machine learning, SARS homolog of this peptide is a known B-cell epitope | (*12*) |
| Ep5 | SARS-CoV-2 | M | 1-24 | MADSNGTITVEELKKLLEQWNLVI | 24 | Predicted B-cell epitope in SARS-Cov-2 identified by bioinformatics to have immunodominant region | (*13*) |
| Ep6 | SARS-CoV-2 | M | 132-151 | PLLESELVIGAVILRGHLRI | 20 | Predicted B-cell epitope in SARS-Cov-2 identified by bioinformatics to have immunodominant region | (*13*) |
| Ep7 | SARS-CoV-2 | M | 97-111 | IASFRLFARTRSMWS | 15 | Predicted B-cell epitope in SARS-CoV-2 identified by structural biology and machine learning, SARS homolog of this peptide is a known B-cell epitope | (*12*) |
| Ep8 | SARS-CoV-2 | N | 41-61 | RPQGLPNNTASWFTALTQHGK | 21 | Predicted B-cell epitope in SARS-Cov-2 identified by bioinformatics to have immunodominant region n | (*13*) |
| Ep9 | SARS-CoV-2 | N | 152-172 | ANNAAIVLQLPQGTTLPKGFY | 21 | Predicted B-cell epitope in SARS-Cov-2 identified by bioinformatics to have immunodominant region | (*13*) |
| Ep10 | SARS-CoV-2 | N | 264-278 | ATKAYNVTQAFGRRG | 15 | Predicted B-cell epitope in SARS-CoV-2 identified by structural biology and machine learning, SARS homolog of this peptide is a known B-cell epitope | (*12*) |
| Ep11 | SARS-CoV-2 | E | 52-66 | VKPSFYVYSRVKNLN | 15 | Predicted B-cell epitope in SARS-CoV-2 identified by structural biology and machine learning, SARS homolog of this peptide is a known B-cell epitope | (*12*) |
| Ep12 | SARS-CoV-2 | S | 524-598 | VCGPKKSTNLVKNKCVNFNFNGLTGTGVLTESNKKFLPFQ<br>QFGRDIADTTDAVRDPQTLEILDITPCSFGGVSVI | 75 | Predicted B-cell epitope in SARS-Cov-2 identified by bioinformatics to have immunodominant region | (*13*) |
| Ep13 | SARS-CoV-2 | S | 601-640 | GTNTSNQVAVLYQDVNCTEVPVAIHADQLTPTWRVYSTGS | 40 | Predicted B-cell epitope in SARS-Cov-2 identified by bioinformatics to have immunodominant region. Based on the D614G variant. | (*13*) |
| Ep13* | SARS-CoV-2 | S | 601-640 | GTNTSNQVAVLYQGVNCTEVPVAIHADQLTPTWRVYSTGS | 40 | Sequence has the mutation D614G, which increases the fitness of SARS-CoV-2 | (25,26,*53*) |

| | | | | | | | |
|---|---|---|---|---|---|---|---|
| Ep14 | SARS-CoV-2 | S | 61-76 | NVTWFHAIHVSGTNGT | 16 | Predicted B-cell epitope in SARS-CoV-2 identified by immuninformatics | (*14*) |
| Ep15 | SARS-CoV-2 | S | 373-390 | SFSTFKCYGVSPTKLNDL | 18 | Predicted B-cell epitope in SARS-CoV-2 identified by immuninformatics | (*14*) |
| Ep16 | SARS-CoV-2 | N | 354-400 | NKHIDAYKTFPPTEPKKDKKKKADETQALPQRQKKQQTVTLLPAADL | 47 | Predicted B-cell epitope in SARS-Cov-2 identified by bioinformatics to have immunodominant region | (*13*) |
| Ep17 | SARS-CoV-2 | S | 319-529 | RVQPTESIVRFPNITNLCPFGEVFNATRFASVYAWNRKRISNCVADYSVLYNSASFSTFKCYGVSPTKLNDLCFTNVYADSFVIRGDEVRQIAPGQTGKIADYNYKLPDDFTGCVIAWNSNNLDSKVGGNYNYLYRLFRKSNLKPFERDISTEIYQAGSTPCNGVEGFNCYFPLQSYGFQPTNGVGYQPYRVVVLSFELLHAPATVCGPKK | 211 | Based on structure of S protein X-ray structure (PBD: 6VXX) | (*23*) |
| Ep18 | SARS-CoV-2 | S | 488-507 | CYFPLQSYGFQPTNGVGYQP | 20 | Based on X-ray structure of S protein (PBD: 6VXX) | (*23*) |
| Ep19 | SARS-CoV-2 | S | 429-448 | FTGCVIAWNSNNLDSKVGGN | 20 | Based on X-ray structure of S protein (PBD: 6VXX) | (*23*) |
| Ep20 | SARS-CoV-2 | S | 448-466 | NYNYLYRLFRKSNLKPFER | 19 | Based on X-ray structure of S protein (PBD: 6VXX) | (*23*) |
| Ep21 | SARS-CoV-2 | S | 467-487 | DISTEIYQAGSTPCNGVEGFN | 21 | Based on X-ray structure of S protein (PBD: 6VXX) | (*23*) |
| sEp9 | SARS | N | 153-173 | NNNAATVLQLPQGTTLPKGFY | 21 | SARS homolog of Ep9 | |
| mEp9 | MERS | N | 141-161 | NNDSAIVTQFAPGTKLPKNFH | 21 | MERS homolog of Ep9 | |
| hEp9 | HKU-1 | N | 166-186 | TTQEAIPTRFPPGTILPQGYY | 21 | HKU-1 homolog of Ep9 | |
| nEp9 | NL63 | N | 119-136 | NQKPLEPKFSIALPPELS | 18 | NL63 homolog of Ep9 | |

*Residue numbering from protein sequences deposited in GenBank. Specifically, the accession numbers were as follows: S protein (YP_009724390.1), M protein (YP_009724393.1), N protein (YP_009724397.2) and E protein (YP_009724392.1) from SARS-CoV-2 and N protein from SARS (NP_828855.1), MERS (YP_009047211.1), HKU-1 (YP_173242.1), or NL63 (YP_003771.1).

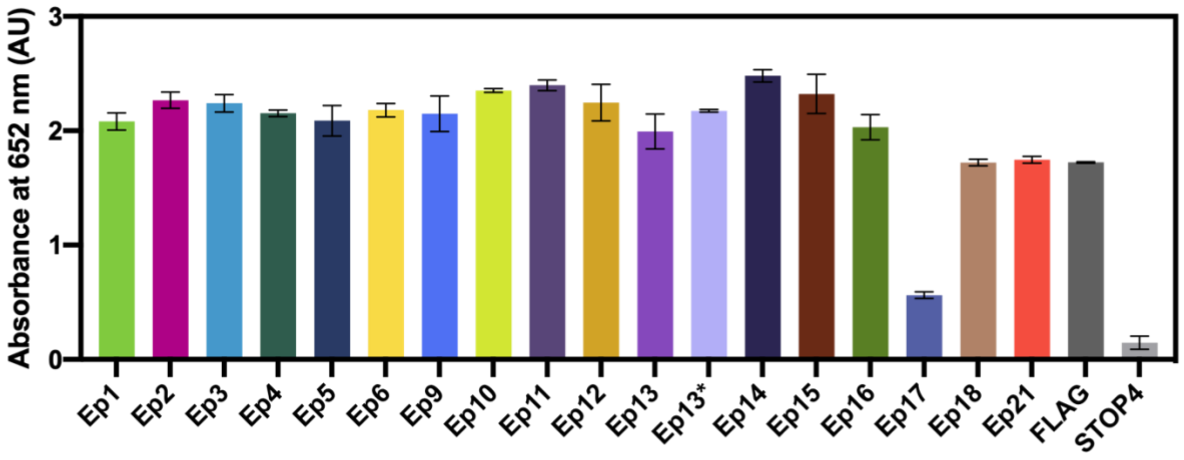

**Fig. S1. Quality control ELISA (QC ELISA) for phage-displayed, epitope candidates.** Anti-FLAG antibodies (1:1000 in coating buffer) were immobilized on a microtiter plate. Subsequent steps followed the ELISA protocol provided here. Error bars represent SEM (n = 3). Ep8 was apparently toxic to *E. coli*, and Ep7 and Ep19 repeatedly failed sequencing quality controls after phage propagation.

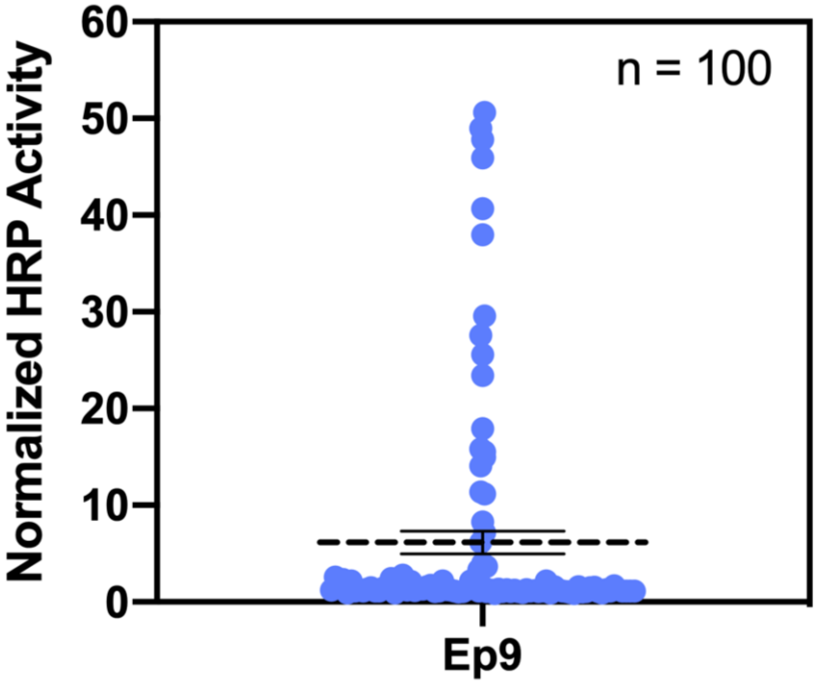

**Fig. S2. Ep9 epitope binding profile of additional COVID-19 patients.** Plasma from 100 COVID-19 patients (1:100) were immobilized on a microtiter plate and binding with phage displayed Ep9 was detected with anti-M13-HRP antibodies. This subset of patients did not have detailed medical records available and thus further analysis of disease progression was not possible. Results demonstrate that 25% of patients test positive for the presence of αEp Abs, as determined using t-test statistical analysis. Error bars represent SEM.

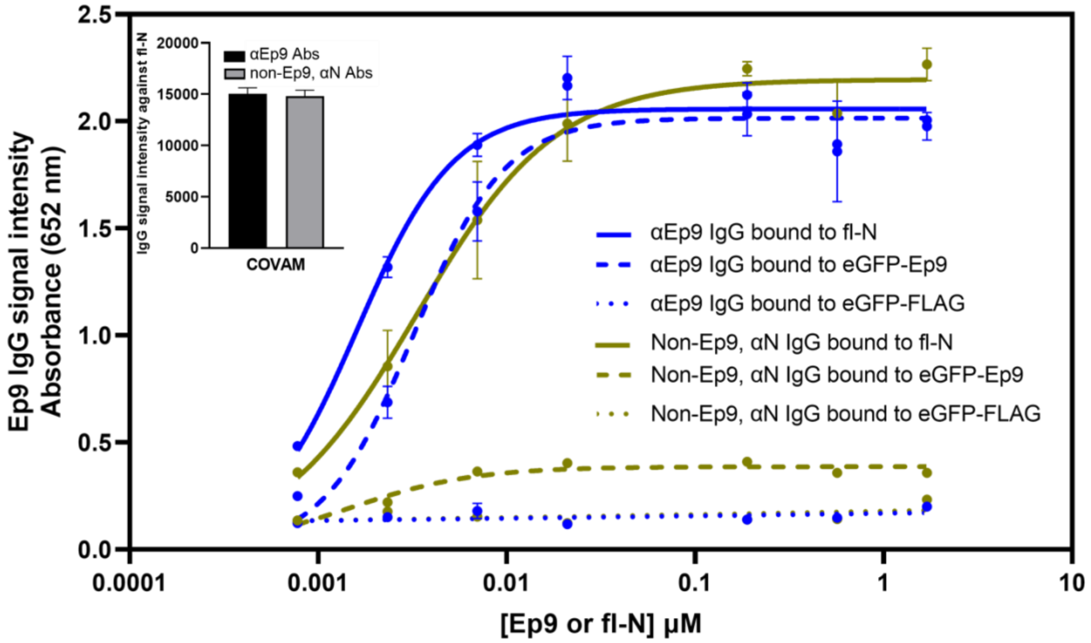

| **Best-fit values (95%CI)** | **αEp9 Abs bound to Ep9** | **αEp9 Abs bound to fl-N** | **αN, non Ep9 Abs bound to fl-N** |
|---|---|---|---|
| **Bmax (@ 652 nm)** | 2.01 (1.89 to 2.14) | 2.06 (1.97 to 2.15) | 2.20 (2.09 to 2.31) |
| **Hill slope** | 1.82 (1.29 to 2.74) | 1.70 (1.30 to 2.25) | 1.17 (0.909 to 1.52) |
| **$EC_{50}$ (nM)** | 3.22 (2.50 to 4.14) | 1.61 (1.33 to 1.95) | 3.33 (2.63 to 4.23) |
| **$R^2$** | 0.960 | 0.967 | 0.958 |

**Fig. S3. Binding of αN IgGs and αEp9 IgGs to Ep9 and full-length N protein**. This ELISA measures dose-dependent binding of αN IgGs from plasma pooled from five αEp9 positive patients and five non-Ep9, αN positive patients to eGFP-Ep9 (dashed line), eGFP negative control (eGFP-FLAG, dotted line) or full-length N protein (fl-N, solid line). Varying doses of Ep9 or fl-N were immobilized on microtiter plates, and binding of pooled patient plasma (1:100) was detected using α-Fc IgG-HRP Abs (1:10,000). Pooled patients were matched by similar αN IgG binding signal in COVAM analysis (inset). Non-linear lines of best-fit for binding saturation are represented. Statistical comparisons of Bmax, Hill slope and $EC_{50}$ between groups, determines that binding of αEp9 IgGs to fl-N or eGFP-Ep9, and non-Ep9, αN IgGs to fl-N are significantly different ($p<0.0001$). Error bars represent ± SD. The data demonstrates that the $EC_{50}$ value of αEp9 Abs is equal to the cumulative $EC_{50}$ of all other αN Abs in patients lacking the αEp9 Abs. In the presence of the αEp9 Abs, the apparent binding levels of αN Abs against fl-N approximately doubles.

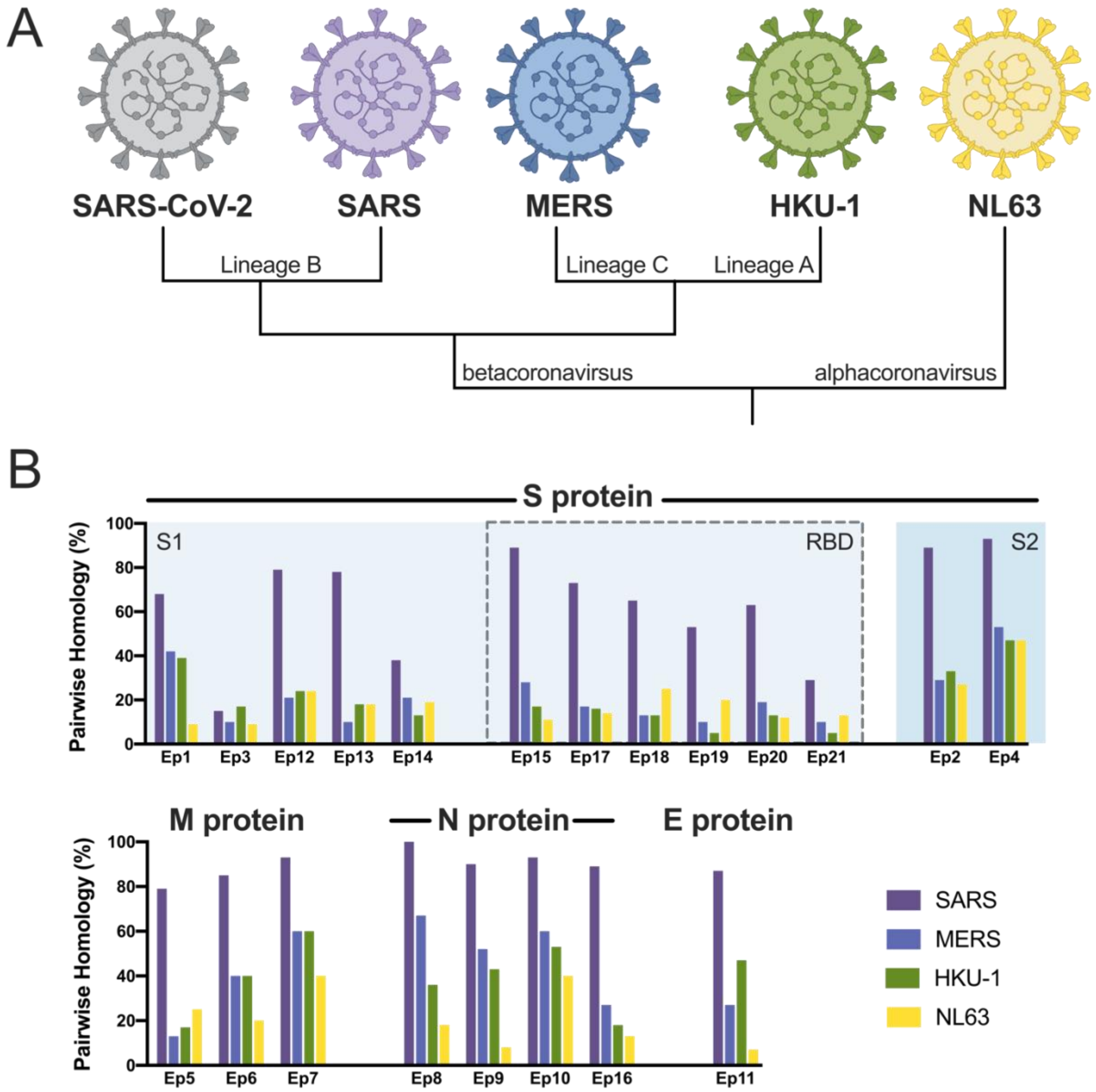

**Fig. S4. Epitope homology of SARS-CoV-2 with four phylogenetically related coronaviruses known to infect humans.** A) Evolutionary lineages of the human coronaviruses investigated here, including the highly pathogenic (SARS-CoV-2, SARS, and MERS) and the less virulent (HKU-1 and NL63). B) The pairwise homology (% amino acid identity) between SARS-CoV-2 and the indicated coronavirus. Labels (top) indicate the proteins and domains (e.g., S1) from which the epitopes are derived.

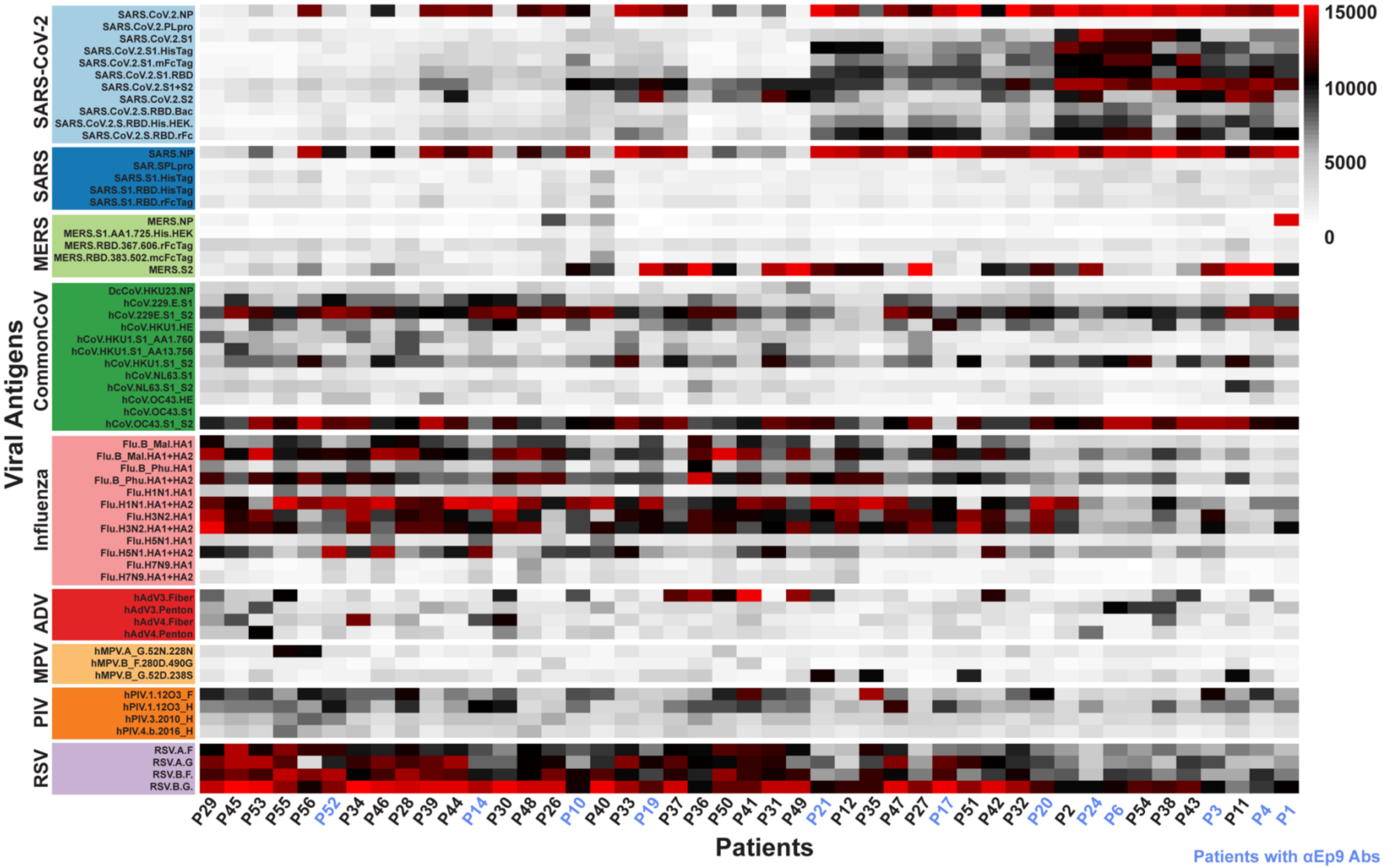

**Fig. S5. COVAM data showing the variation in IgG seroreactivity of patient plasma.** The heatmap shows normalized signal intensity from plasma samples (n = 45). Plasma samples are in columns and sorted left to right by increasing average intensity to differentially reactive IgG, and viruses are in rows sorted by decreasing average seroreactivity.

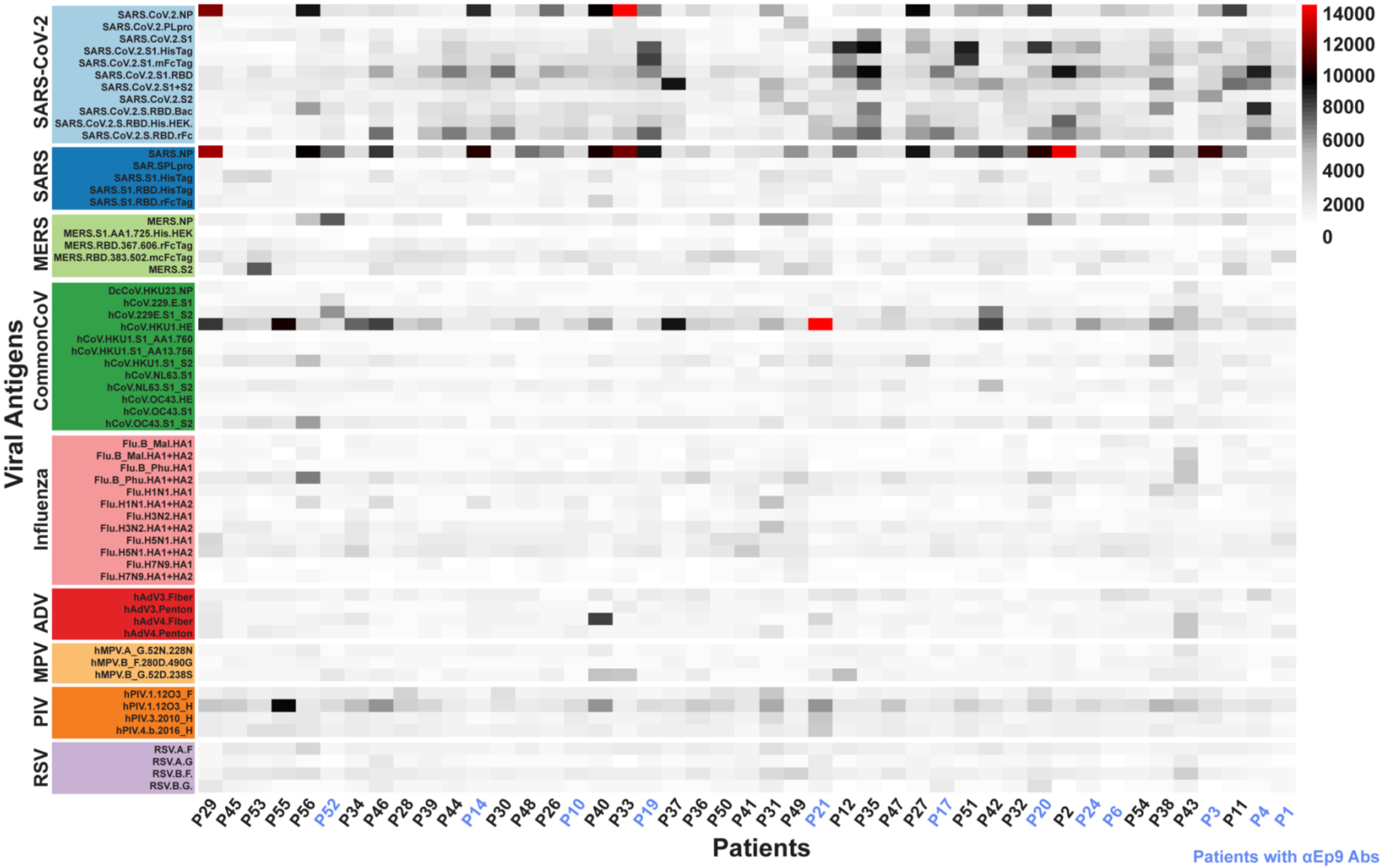

**Fig. S6. Variation in IgM seroreactivity of patient plasma.** Heatmap showing normalized signal intensity from plasma samples (n = 45). Plasma samples are in columns and sorted left to right by increasing average intensity to differentially reactive IgM, and viruses are in rows sorted by decreasing average seroreactivity.

**Table S2. Demographics and clinical characteristics of COVID-19 patients categorized by αEp9 Abs response.**

| Characteristics | No αEp9 Abs (n=63) | αEp9 Abs (n=23) | p-value |
|---|---|---|---|
| **Demographics** | | | |
| Age (± SD) | 49.75 (± 18.45) | 47.26 (± 18.45) | 0.5668 |
| Gender F: M (%) | 21:42 (44.4/66.7) | 10:13 (43.5/56.5) | 0.4502 |
| Ethnicity n, (%) (Hispanic: Asian: Caucasian: Black: Other) | 15 (65.2): 4 (17.4): 3 (13.0): 1 (4.3): 0 (0) | 39 (61.9): 8 (12.7): 9 (14.3) :3 (4.8) :4 (6.3) | 0.7760 |
| BMI (± SD) | 28.89 (± 6.445) | 32.06 (± 7.896) | 0.0642 |
| **Preconditions, n (%)** | | | |
| Hypertension | 23 (36.5) | 10 (43.5) | 0.6203 |
| Diabetes | 21 (33.3) | 6 (26.1) | 0.6065 |
| CVD | 6 (9.5) | 2 (8.7) | 1.0000 |
| CAD | 6 (9.5) | 2 (8.7) | 1.0000 |
| CKD/ESRD | 6 (9.5) | 2 (8.7) | 1.0000 |
| Asthma/COPD | 8 (12.7) | 3 (13.0) | 1.0000 |
| Obesity | 24 (38.1) | 13 (56.5) | 0.1461 |
| Cancer | 2 (3.17) | 3 (13.0) | 0.1163 |
| **Symptoms, n (%)** | | | |
| Total Days of Symptoms | 9.8 (± 8.98) | 17 (± 10.13) | 0.0059** |
| Cough | 43 (68.3) | 15 (65.2) | 0.7997 |
| Dyspnea/SOB | 28 (44.4) | 11 (47.8) | 0.8108 |
| Myalgia/Fatigue | 17 (27.0) | 8 (34.8) | 0.5926 |
| Headache | 12 (19.0) | 2 (8.7) | 0.3349 |
| Chest pain | 7 (11.1) | 3 (13.0) | 1.0000 |
| Anosmia | 4 (6.3) | 2 (8.7) | 0.6561 |
| Stroke-like Symptoms | 0 | 2 (8.7) | 0.0692 |
| Abdominal pain | 3 (4.8) | 0 | 0.5611 |
| Pulmonary symptoms^ (Pneumonia: Other: None) | 16 (25.4): 36 (52.38): 8 (12.7) | 13 (56.5): 7 (30.4): 1 (4.3) | 0.0142* |
| **Severity, n (%)** | | | |
| Asymptomatic | 3 | 0 | 0.5611 |
| Non-severe: Severe^^ | 51:12 (n, severity 19.0%) | 10:13 (n, severity 56.5%) | 0.0013** |
| Days in Hospital | 5.79 (± 8.01) | 10.95 (± 10.74) | 0.0183* |
| Days in ICU | 12.63 (± 13.19) n=11 | 12.50 (± 6.93), n=12 | 0.8004 |
| Days on ventilator | 14.00(± 3.96), n=6 | 12.86 (± 5.40), n=7 | 0.7934 |

Results are presented as mean ± standard deviation (SD) or patient number (n) and percentage of population (%). P-values for continuous variables are calculated using unpaired, two-tailed T-tests. P-values for categorical variables use Fisher's exact test for single value parameters, and Chi-squared test for multi-group variables. *****, ****** p-values < 0.05, 0.01, respectively.

**^** Pulmonary symptoms are based descriptive reports of X-ray and CT scans. "Other" pulmonary symptoms include, but are not limited to, atelectasis, pleural scarring, pleural effusion, pulmonary edema, mild peribronchial thickening.
**^^** non-severe include ER and In-patients only, severe includes patients in the ICU, on the ventilator or death.

BMI = body mass index, CVD = cardiovascular disease, CAD = coronary artery disease, CKD = chronic kidney disease, ESRD = end-stage renal disease, SOB = shortness of breath, COPD = chronic obstructive pulmonary disease

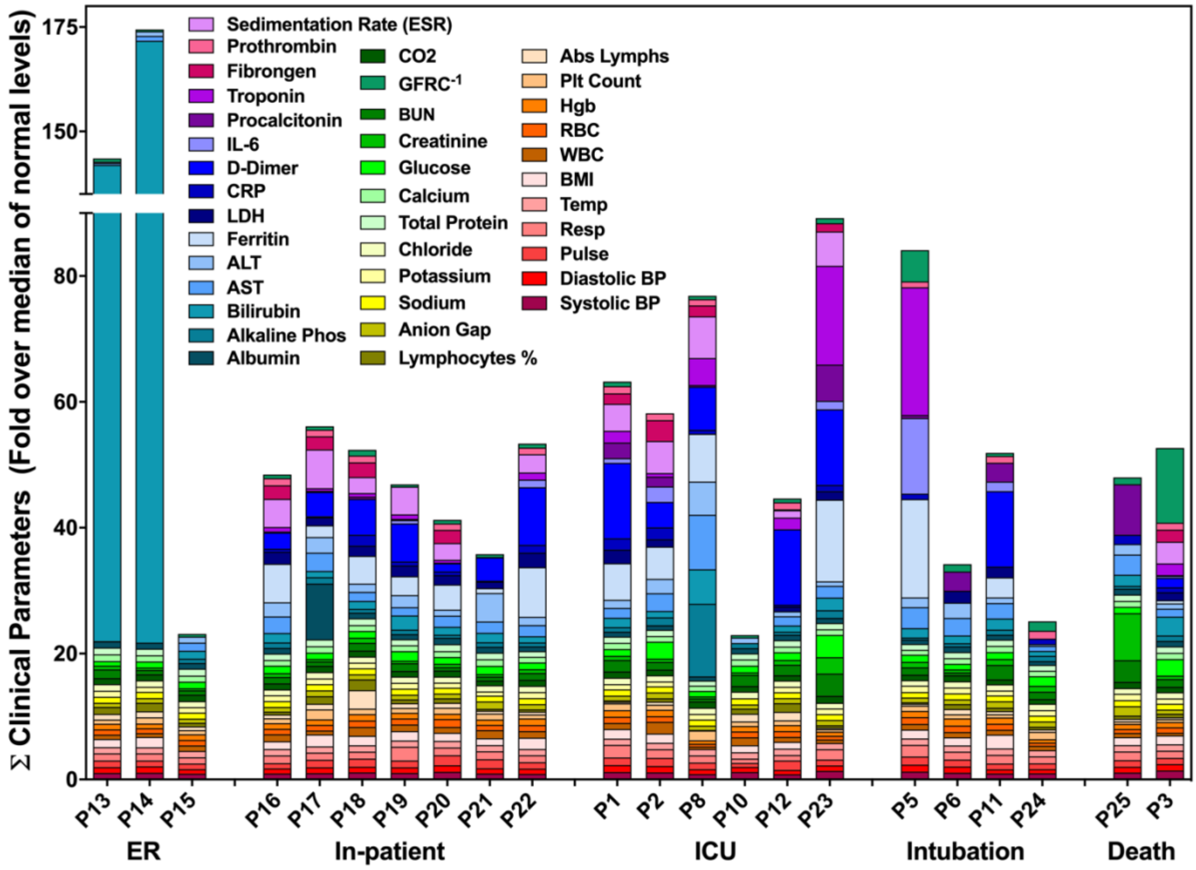

**Fig. S7. Comparison of disease severity and clinical parameters of patients with αEp9 Abs.** The data shown represents the fold change of each clinical parameter over the mean of the normal range. The sum of all the fold changes of the clinical parameters for each Ep9-responsive patient is binned according to COVID-19 disease severity. For facile visualization and comparison of clinical biomarkers between Ep9-reponsive patients, the values of each parameter were normalized to fold over the mean of healthy values. No significant trends in clinical parameters (color indicated) were observed with increased disease severity or relative to patients lacking αEp9 Abs.

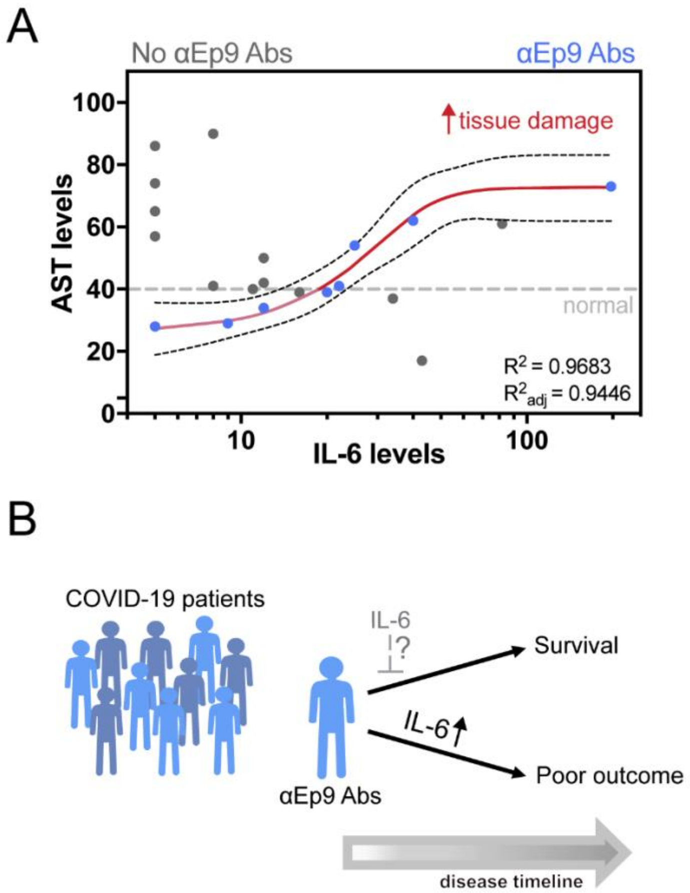

**Fig. S8. Association of COVID-19 patients having αEp9 Abs with inflammatory cytokine and tissue damage markers. A**) Association between the inflammatory cytokine, IL-6, and the tissue damage marker, aspartate transaminase (AST), shows a sigmoidal curve fit for patients with αEp9 Abs, $R^2 = 0.9683$, Spearman's correlation coefficient = 1.0, $p < 0.0001$. **B**) Schematic of patients with αEp9 Abs with increasing IL-6 levels leading to poor outcomes. We hypothesize patients with αEp9 Abs could benefit from IL-6 inhibition early in the disease, such as monoclonal antibody drugs targeting IL-6 or its receptor (IL6R), to disrupt a cytokine storm and reduce severe outcomes.

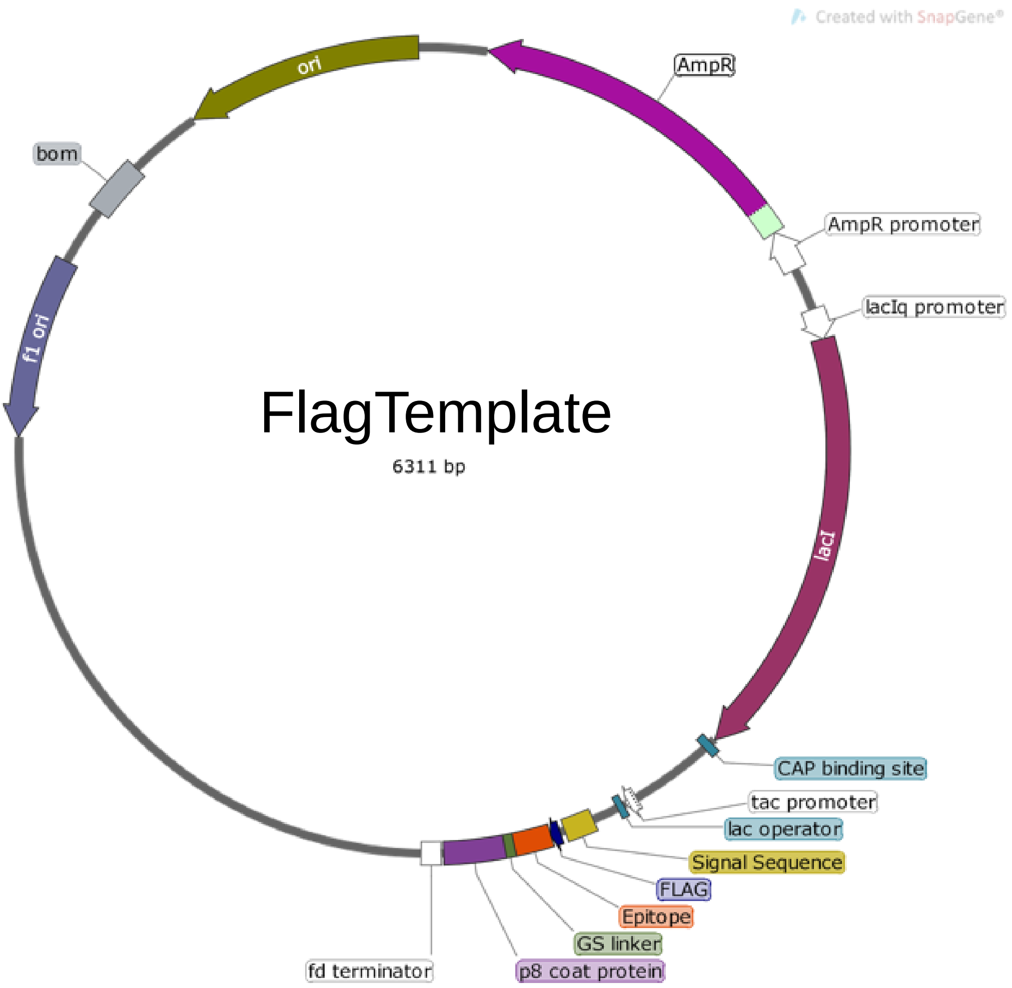

**Fig. S9. Schematic of the FlagTemplate phagemid used for cloning phage-displayed epitopes.** The phagemid, termed FlagTemplate, for the subcloning of SARS-CoV-2, SARS, HKU-1 and NL63 epitopes encodes an N-terminal FLAG tag, followed by a GSG linker to the epitope before a C-terminal GGGSGSSS linker to the P8 coat protein of M13-phage.

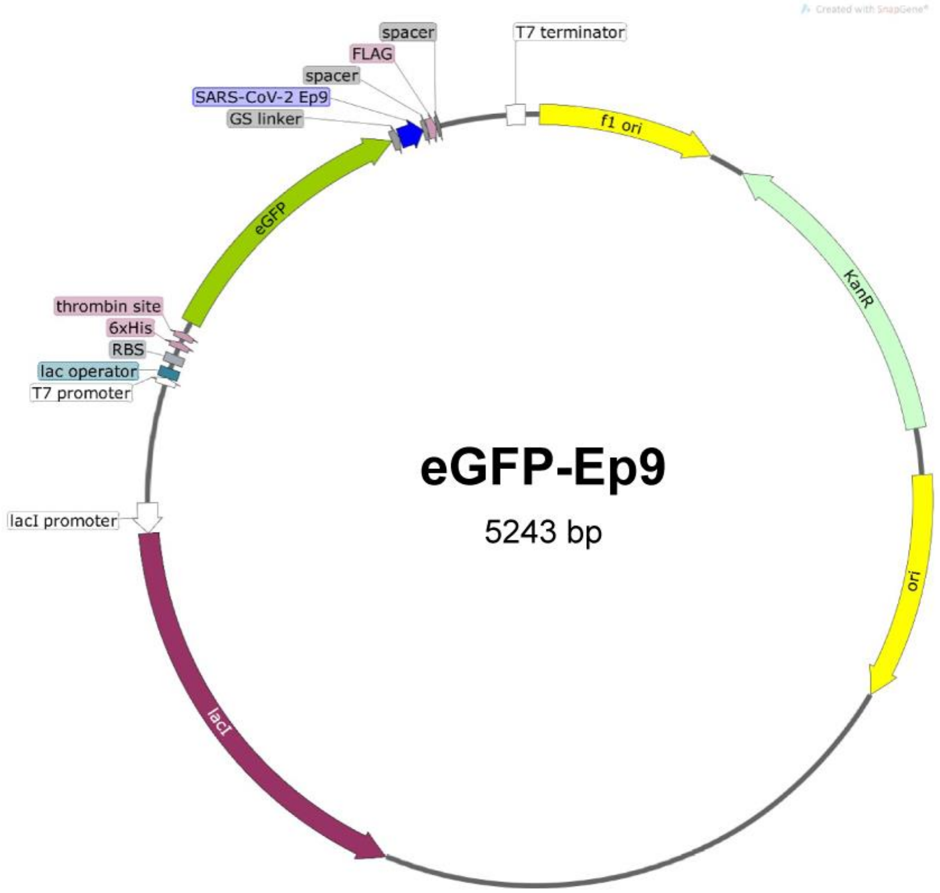

**Fig. S10. The plasmid map for recombinant expression of the eGFP-Ep9 fusion.** Ep9 was subcloned into a pET28-eGFP fusion vector with a C-terminal FLAG tag. The fusion protein is connected through a linker (GGGSGSS), and two spacers flank the N- and C-termini of the FLAG-tag (SGSG and GSG, respectively). The plasmid backbone lacking the Ep9 sequence was used to express the eGFP-FLAG negative control.

**Table S3. Oligos used for cloning of phage-displayed putative epitope for SARS-CoV-2 and ortholog Ep9 epitope from SARS, MERS, HKU-1, and NL63.**

| Oligo# | Sequence (5' to 3') with insertions denoted in lowercase and substitutions in bold | Product | Mutagenesis | Rounds |
|---|---|---|---|---|
| Oligo1 | aggaagtggaggtggaggatccgggagctccagcCCGAGGGTGACGATCCCG | FlagTemplate | Q5 | 1 |
| Oligo2 | ttatcatcgtcatctttataatcaaccaatgcataGCCGAGGCGGAAAACATC | | | |
| Oligo3 | attaaagtcgttcaccgtcgaaaaggaatctatcagacctctaacGGTGGAGGATCCGGGAGC | Ep1 | Q5 | 1 |
| Oligo4 | gtacactttgtttcactcagtggatctaatgcacaatcgaccgcatcTTTATCATCGTCATCTTTATAATCAAC CAATGC | | | |
| Oligo5 | gagcaagccttctaagcgctctttcattgaaGGTGGAGGATCCGGGAGC | Ep2 | Q5 | 1 |
| Oligo6 | gggtcaggcaggatctgcgagaatccacttccTTTATCATCGTCATCTTTATAATCAACCAATGC | | | |
| Oligo7 | tacacagttacctcccgcgtatacaaatGGTGGAGGATCCGGGAGC | Ep3 | Q5 | 1 |
| Oligo8 | cgagttgtcaagttcacacatccacttccTTTATCATCGTCATCTTTATAATCAACCAATGC | | | |
| Oligo9 | ttcttgcacgtgacgtatgtgcctgctGGTGGAGGATCCGGGAGC | Ep4 | Q5 | 1 |
| Oligo10 | caccactccatggggcgctccacttccTTTATCATCGTCATCTTTATAATCAACCAATGC | | | |
| Oligo11 | ggagctgaaaaaactgttggaacaatggaaccttgtaatcGGTGGAGGATCCGGGAGC | Ep5 | Q5 | 1 |
| Oligo12 | tctacggtaatcgtaccgttcgagtccgccattccacttccTTTATCATCGTCATCTTTATAATCAACCAATGC | | | |
| Oligo13 | tggagctgtgatcttacgcggacacctgcgtatcGGTGGAGGATCCGGGAGC | Ep6 | Q5 | 1 |
| Oligo14 | atcactaattctgattccaacaggggtccacttccTTTATCATCGTCATCTTTATAATCAACCAATGC | | | |
| Oligo15 | ttcgcacgcactcgctccatgtggtctGGTGGAGGATCCGGGAGC | Ep7 | Q5 | 1 |
| Oligo16 | caagcggaagctcgcaattccacttccTTTATCATCGTCATCTTTATAATCAACCAATGC | | | |
| Oligo17 | gcttcgtggttcactgcgcttacccagcacggaaagGGTGGAGGATCCGGGAGC | Ep8 | Q5 | 1 |
| Oligo18 | tgtattattaggcagcccttgagggcgtccacttccTTTATCATCGTCATCTTTATAATCAACCAATGC | | | |
| Oligo19 | tcaagggactaccttgcccaaggggttctatGGTGGAGGATCCGGGAGC | Ep9 | Q5 | 1 |
| Oligo20 | ggtaattgtaacacgattgcagcgttattagcTCCACTTCCTTTATCATCGTCATCTTTATAATC | | | |
| Oligo21 | gtaacccaagcgttcggtcgccgcgggGGTGGAGGATCCGGGAGC | Ep10 | Q5 | 1 |
| Oligo22 | attatacgcttttgtagctccacttccTTTATCATCGTCATCTTTATAATCAACCAATGC | | | |
| Oligo23 | gtctactctcgtgtaaaaaacttgaatGGTGGAGGATCCGGGAGC | Ep11 | Q5 | 1 |
| Oligo24 | gtaaaaggaaggcttcactccacttccTTTATCATCGTCATCTTTATAATCAACCAATGC | | | |
| Oligo25 | tttcaactttaatggcctgacggggaccggagtcctgactgaatccaatGGTGGAGGATCCGGGAGC | Ep12 | Q5 | 3 |
| Oligo26 | ttgacacacttatttttaaccaggtttgttgacttcttcggcccgcatacTTTATCATCGTCATCTTTATAATC AACCAATGC | | | |
| Oligo27 | agacgctgttcgtgacccacagactctggagattttggacattacacctGGTGGAGGATCCGGGAGC | | | |
| Oligo28 | gttgtatcggcgatgtcgcgtccaaattgctggaacggcagaaatttttATTGGATTCAGTCAGGACTCCG | | | |
| Oligo29 | tgtctccgtcatcGGTGGAGGATCCGGGAGC | | | |
| Oligo30 | cctccgaatgaacaAGGTGTAATGTCCAAAATCTCCAGAG | | | |
| Oligo31 | ctgtatcaggatgtcaattgcacagaagtcGGTGGAGGATCCGGGAGC | Ep13 | Q5 | 2 |
| Oligo32 | tacagcaacttggttacttgtgtttgtcccTTTATCATCGTCATCTTTATAATCAACCAATGC | | | |
| Oligo33 | cccacttggcgcgtctacagcacaggcagtGGTGGAGGATCCGGGAGC | | | |
| Oligo34 | cgtcagttggtcggcgtggatggctaccggGACTTCTGTGCAATTGACATCCTGATAC | | | |
| Oligo35 | GCTGTACCAG**ggc**GTGAACTGTA | Ep13* | Q5 | 1 |
| Oligo36 | ACTGCCACCTGATTGCTG | | | |
| Oligo37 | tatccacgtatcgggtacgaatggaacgGGTGGAGGATCCGGGAGC | Ep14 | Q5 | 1 |
| Oligo38 | gcgtggaaccacgtcacatttccacttccTTTATCATCGTCATCTTTATAATCAACCAATGC | | | |
| Oligo39 | tggagtgtctcccactaaattgaacgaccttGGTGGAGGATCCGGGAGC | Ep15 | Q5 | 1 |
| Oligo40 | tagcatttgaaggttgagaacgatccacttccTTTATCATCGTCATCTTTATAATCAACCAATGC | | | |
| Oligo41 | tcccccgacggagccgaagaaggataagaaaaaaaaaGGTGGAGGATCCGGGAGC | Ep16 | Q5 | 2 |
| Oligo42 | aaggttttatacgcatcaatgtgtttatttccacttccTTTATCATCGTCATCTTTATAATCAACCAATGC | | | |
| Oligo43 | acttccacagggaacgacactgccaaagggatttGGTGGAGGATCCGGGAGC | | | |
| Oligo44 | tgcagtacagtggcagcattgttatttccacttccTTTATCATCGTCATCTTTATAATCAACCAATGC | | | |
| Oligo45 | GGTGGAGGATCCGGGAGCTCCAGC | GA Vector | - | - |
| Oligo46 | TTTATCATCGTCATCTTTATAATCAACCAATGCATAAGCCGAGGC | | | |
| Oligo47 | CTGTATGTGGGCCCAAAAAGGGTGGAGGATCCGGGAGCTC | Ep17 | GA | |
| Oligo48 | GGCTGTACGCGTCCACTTCCTTTATCATCGTCATCTTTATAATCAACCAATGCATAAG | | | |
| Oligo49 | ACGATGATAAAGGAAGTGGACGCGTACAGCCCACTGAAAG | | | |

| | | | | |
|---|---|---|---|---|
| Oligo50 | GAGCTCCCGGATCCTCCACCCTTTTTGGGCCCACATACAGTCG | | | |
| Oligo51 | caaccgacaaatggtgtgggttaccagccgGGTGGAGGATCCGGGAGC | Ep18 | Q5 | 1 |
| Oligo52 | gaaaccatatgactgcaagggaaaataacaTCCACTTCCTTTATCATCGTCATCTTTATAATC | | | |
| Oligo53 | taataacttggattccaaagtaggaggcGGTGGAGGATCCGGGAGC | Ep19 | Q5 | 1 |
| Oligo54 | gaattccaggcaataacacacccagtgaaTCCACTTCCTTTATCATCGTCATCTTTATAATC | | | |
| Oligo55 | gaaatctaacttgaaaccgtttgaaaggGGTGGAGGATCCGGGAGC | Ep20 | Q5 | 1 |
| Oligo56 | cggaaaagcctgtaaagatagttgtaattTCCACTTCCTTTATCATCGTCATCTTTATAATC | | | |
| Oligo57 | cacgccttgcaatggggtagagggttttaatGGTGGAGGATCCGGGAGC | Ep21 | Q5 | 1 |
| Oligo58 | gaccctgcctggtagatctccgttgagatgtcTCCACTTCCTTTATCATCGTCATCTTTATAATC | | | |
| Oligo59 | caaccgacaaatggtgtgggttaccagccgGGTGGAGGATCCGGGAGC | Ep22 | Q5 | 1 |
| Oligo60 | gaaaccatatgactgcaagggaaaataacaTCCACTTCCTTTATCATCGTCATCTTTATAATC | | | |
| Oligo61 | acttccacagggaacgacactgccaaagggatttGGTGGAGGATCCGGGAGC | sEp9 | Q5 | 1 |
| Oligo62 | tgcagtacagtggcagcattgttatttccacttccTTTATCATCGTCATCTTTATAATCAACCAATGC | | | |
| Oligo63 | tccgggtacaaagttaccaaagaacttccacGGTGGAGGATCCGGGAGC | mEp9 | Q5 | 1 |
| Oligo64 | gcaaattgagtaactatcgctgaatcattgttTCCACTTCCTTTATCATCGTCATCTTTATAATC | | | |
| Oligo65 | tcccggaactattttaccccaaggatactatGGTGGAGGATCCGGGAGC | hEp9 | Q5 | 1 |
| Oligo66 | gggaatctagtgggaatcgcctcctgagtagtTCCACTTCCTTTATCATCGTCATCTTTATAATC | | | |
| Oligo67 | agtattgccttgccacctgagttatctGGTGGAGGATCCGGGAGC | nEp9 | Q5 | 1 |
| Oligo68 | aaatttcggttcaagcggcttttgattTCCACTTCCTTTATCATCGTCATCTTTATAATC | | | |
| Oligo69 | TTTTGCGCCGACATCATAACGGTTCT | pm1165a_P8 | - | - |
| Oligo70 | TATGGGGTTTTGCTAAACAACTTTCAACAG | | | |
| Oligo71 | agatgacgatgataaaggaagtggataataaGGCGGTTCGGCAGGCGAA | eGFP-FLAG | Q5 | 1 |
| Oligo72 | ttataatcgctggagctcccggatcctccaccTTTGTATAGTTCATCCATGCCATGTGTAATCCC | | | |
| Oligo73 | tcaagggactaccttgcccaaggggttctatAGCGGAAGTGGAGATTATAAAGATGAC | eGFP-Ep9 | Q5 | 1 |
| Oligo74 | ggtaattgtaacacgattgcagcgttattagcGGAGCTCCCGGATCCTCC | | | |
| Oligo75 | TAATACGACTCACTATAGGG | T7 | - | - |
| Oligo76 | GCTAGTTATTGCTCAGCGG | T7 term | - | - |

List of oligos used in Q5 site-directed mutagenesis (Q5) or Gibson Assembly (GA) for the cloning of putative epitopes Ep1-22 for phage display or eGFP recombinant protein expression. For epitope constructs between 100-210 bp in length, multiple rounds of Q5 were denoted.

**Table. S4. Antigens used in the COVAM experiment.**

| Virus | Subtype | Strain | Protein | Gen Bank | Expression | Construct | Source | Cat. # |
|---|---|---|---|---|---|---|---|---|
| **CoV** | Beta | SARS-CoV-2 | N | | Baculovirus | N-(AA)-His-C | Sino | 40588-V08B |
| **CoV** | Beta | SARS-CoV-2 | S1-RBD | | HEK293 | N-(AA)-mFc-C | Sino | 40592-V05H |
| **CoV** | Beta | SARS-CoV-2 | S1 | | HEK293 | N-(AA)-His-C | Sino | 40591-V08H |
| **CoV** | Beta | SARS-CoV-2 | S1 | | HEK293 | N-(AA)-mFc-C | Sino | 40591-V02H |
| **CoV** | Beta | SARS-CoV-2 | S1 | | HEK293 | N-(AA)-Fc-C | Sino | 40591-V05H1 |
| **CoV** | Beta | SARS-CoV-2 | S2 | | Baculovirus | N-(AA)-His-C | Sino | 40590-V08B |
| **CoV** | Beta | SARS-CoV-2 | S1+S2 | | Baculovirus | N-(AA)-His-C | Sino | 40589-V08B1 |
| **CoV** | Beta | SARS | PLpro | AAX16193.1 | E. coli | N-(AA1541-1859)-His-C | Sino | 40524-V08E |
| **CoV** | Beta | SARS | S1-RBD | AAX16192.1 | Baculovirus | N-(AA306-527)-Fc-C | Sino | 40150-V31B2 |
| **CoV** | Beta | SARS | S1-RBD | AAX16192.1 | Baculovirus | N-(AA306-527)-His-C | Sino | 40150-V08B2 |
| **CoV** | Beta | SARS | S1 | AAX16192.1 | Baculovirus | N-(AA1-667)-His-C | Sino | 40150-V08B1 |
| **CoV** | Beta | SARS | N | NP_828858.1 | Baculovirus | N-(AA1-422)-His-C | Sino | 40143-V08B |
| **CoV** | Beta | MERS | N | AFS88943.1 | Baculovirus | N-(AA1-413)-His-C | Sino | 40068-V08B |
| **CoV** | Beta | MERS | S1-RBD | AFS88936.1 | Baculovirus | N-(AA383-502)-Fc-C | Sino | 40071-V05B |
| **CoV** | Beta | MERS | S1-RBD | AFS88936.1 | Baculovirus | N-(AA383-502)-rFc-C | Sino | 40071-V31B |
| **CoV** | Beta | MERS | S1-RBD | AFS88936.1 | Baculovirus | N-(AA367-606)-rFc-C | Sino | 40071-V31B1 |
| **CoV** | Beta | MERS | S1-RBD | AFS88936.1 | Baculovirus | N-(AA367-606)-His-C | Sino | 40071-V08B1 |
| **CoV** | Beta | MERS | S1 | AFS88936.1 | HEK293 | N-(AA1-725)-His-C | Sino | 40069-V08H |
| **CoV** | Beta | MERS | S1 | AFS88936.1 | Baculovirus | N-(AA1-725)-His-C | Sino | 40069-V08B1 |
| **CoV** | Beta | MERS | S1+S2 | AFS88936.1 | Baculovirus | N-(AA1-1297)-His-C | Sino | 40069-V08B |
| **CoV** | Beta | MERS | S2 | AFS88936.1 | Baculovirus | N-(AA726-1296)-His-C | Sino | 40070-V08B |
| **CoV** | Alpha | NL63 | S1 | A0A1L2YVI8 | HEK293 | N-(AA19-717)-His-C | Sino | 40600-V08H |
| **CoV** | Alpha | NL63 | S1+S2 | A0A1L2YVI8 | Baculovirus | N-(AA19-1296)-His-C | Sino | 40604-V08B |
| **CoV** | Alpha | 229E | S1 | A0A1L7B942 | HEK293 | N-(AA16-536)-His-C | Sino | 40601-v08H |
| **CoV** | Alpha | 229E | S1+S2 | A0A1L7B942 | Baculovirus | N-(AA16-1115)-His-C | Sino | 40605-V08B |
| **CoV** | Beta | HKU-1 | S1 | YP_173238.1 | HEK293 | N-(AA1-760)-His-C | Sino | 40021-V08H |
| **CoV** | Beta | HKU-1 | S1 | Q0ZME7 | HEK293 | N-(AA13-756)-His-C | Sino | 40602-V08H |
| **CoV** | Beta | HKU-1 | S1+S2 | Q0ZME7 | Baculovirus | N-(AA13-1295)-His-C | Sino | 40606-V08B |
| **CoV** | Beta | HKU-1 | HE | Q0ZME7 | HEK293 | N-(AA16-394)-His-C | Sino | Custom |
| **CoV** | Beta | HKU23-368F | N | AHN64796.1 | HEK293 | N-(AA1-448)-His-C | Sino | 40458-V08B |
| **CoV** | Beta | OC43 | S1 | AVR40344.1 | HEK293 | N-(AA13-533)-His-C | Sino | Custom |
| **CoV** | Beta | OC43 | S1+S2 | AVR40344.1 | Baculovirus | N-(AA13-1304)-His-C | Sino | 40607-V08B |

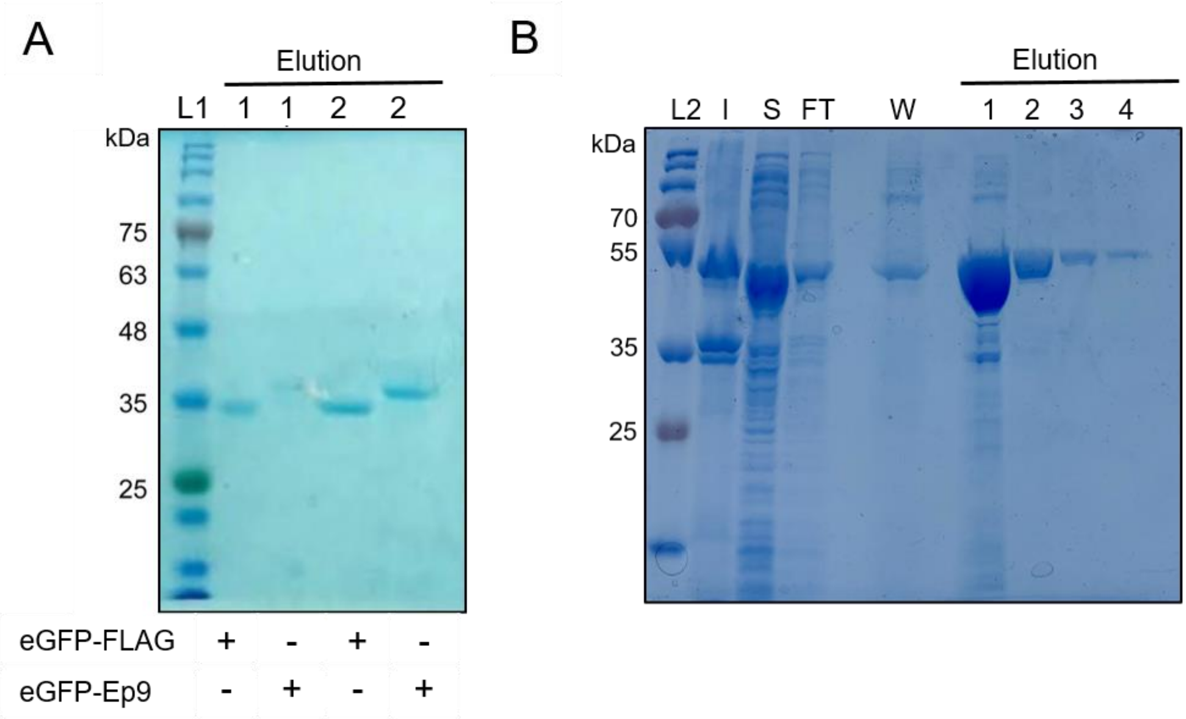

**Fig. S11. eGFP-FLAG, eGFP-Ep9 and full-length N protein purity assessed by 10% SDS-PAGE.** Representative SDS-PAGE to visualize **A)** eGFP-FLAG and eGFP-Ep9 or **B)** full-length N protein after immobilized metal ($Ni^{2+}$) affinity chromatography purification. Elution fraction #2 of eGFP-FLAG, eGPF-Ep9, and elution fraction #3 were purified to >95% homogeneity, and migrated at the expected mass of ≈32, ≈34, and ≈48 kDa, respectively. Elution fraction #1 of eGFP-FLAG and eGFP-Ep9 was diluted 20-fold. L1, BLUE stain (Goldbio) and L2, Prestain PAGE-Ruler Plus (Thermo Fisher Scientific) protein ladder was used as reference. I = insoluble fraction, S = soluble fractions, FT = flow-through, W = wash.
